# mRNA concentration–dependent translation enables rapid and sharp patterning in resource-constrained *Drosophila* embryos

**DOI:** 10.64898/2026.06.15.732008

**Authors:** Jiayi Chen, Xinru Wang, Yue Sun, Bingxiang Xu, Feng Liu

## Abstract

One of the fundamental challenges in protein expression during embryogenesis is the establishment of dynamic protein patterns under the resource constraint. This constraint is particularly stringent in enclosed *Drosophila* early embryogenesis, where protein synthesis relies exclusively on maternally supplied resources, yet its zygotic protein patterns still exhibit sharp boundaries and dynamic abundance changes for body plan development. Here, by integrating single-molecule imaging with mathematical modeling, we reveal that an mRNA concentration– dependent translation enables sharp and dynamics patterning even under resource constraint in embryos. Firstly, we reveal pronounced spatial heterogeneity in the fraction of *hunchback* (*hb*) mRNAs engaged in translation, challenging the conventional assumption of uniform translation rates. We further demonstrate that *hb* mRNA concentration is a dominant predictor of this heterogeneity, showing an increasing-saturation relationship between this translating fraction and mRNA abundance. This regulatory strategy enables combination of sharp Hb protein boundaries and close tracking of protein level to mRNA concentration decline under slow protein turnover rates, correlated to the resource efficiency in embryos. Together, our findings identify a translational control strategy that links molecular-scale regulation to embryonic pattern formation under resource constraints, and highlight mRNA concentration–dependent translation as an effective way simultaneously enabling dynamic and economical spatial gene expression in biological systems.

## INTRODUCTION

Gene expression can often be spatially and temporally dynamic, enabling cells to respond to changes in either internal state or external cues, especially in multicellular systems. Meanwhile, the resource burden of gene expression imposes a fundamental constraint that can lead to a trade-off between resource efficiency and gene expression characteristics^1^. For example, increasing molecular degradation rates can accelerate system pace^2^, or has been theoretically proposed to sharpen a gradient^3^, but at the expense of elevated resource expenditure due to increased molecular turnover.

Despite this constraint, *Drosophila* embryogenesis still achieves both responsiveness and resource efficiency. From macroscopic perspective, the pace of *Drosophila* embryos from fertilized eggs to gastrula is the fastest one compared with multiple model organisms including zebrafish, *Xenopus* and mouse embryogenesis^4^. From microscopic perspective, both the mRNA and protein of zygotic patterning–related genes undergo dynamic expression processes even within the temporal resolution of minutes, accompanied with limited sequential delay between mRNA and protein^5,6^. Moreover, representative developmental pattern Hunchback (Hb) build-ups sharp boundaries along the anterior-posterior (AP) axis of the embryo^7^. If such rapid responsiveness and boundary sharpening were achieved primarily through increased protein degradation, the embryo would incur a substantial resource cost. However, the energy supply provided by the enclosed yolk is strictly limited^8^, raising the intriguing question of how *Drosophila* embryos reconcile dynamic pattern formation with resource efficiency.

To understand this paradox, translational regulation could emerge as a compelling control mechanism. Firstly, translation is suggested to constitute the primary energetic investment in protein expression^9^, and its modulation therefore most directly influences the cost of gene expression. In contrast, while protein degradation determines protein lifetime, it represents a comparatively smaller energetic burden^9^. Secondly, translation rates exhibit two orders greater variability across genes than protein degradation rates, further highlighting its capacity for dynamic and context-specific control^1^. Especially, this flexibility is critical in multicellular systems, where spatial heterogeneity of translation distinguish cell types and correlating with compartmentalized functions among different spatial regions, or serves as a potential determinator for cell plasticity and cell fate commitment during development^10,11^. The heterogeneity of translation in multicellular systems requires the spatial-temporal coordination from single-molecule kinetics to the multicellular level orchestration, while the detailed mechanism has yet to be explored. Finally, translation does not operate in isolation but actively communicates with transcription and mRNA stability^12–16^, facilitates more complex regulation in gene expression. Focusing on translation therefore targets a central, tunable node in the gene expression network that can balance speed and resource economy, and shape developmental patterns.

However, a major obstacle to testing this idea is the difficulty of faithfully quantifying translation signal especially in multicellular systems. For instance, the single-molecule translating RNA imaging by coat protein knock-off (TRICK) only detects the first round of translation^17^. Ribosome-bound mRNA mapping (RIBOmap) detects mRNAs with ribosome bindings rather than the exact translation kinetics^10^. The SunTag system does comprehensively quantify translational kinetics^18–23^. However, recent studies point out the distorted measurement results due to the insufficient supply of corresponding detectors. This detector supply issues and aggregation problem^23–25^ can be alleviated by optimizing tagging fluorescent protein of detector, or replacing scFv detector with antibody.

In this work, we reveal that the mRNA concentration–dependent translation of *hb* mRNA relaxes the trade-off between dynamic protein patterning and resource efficiency during *Drosophila* embryogenesis. Hb is a key transcription factor for segmentation determination before and during the onset of zygotic genome activation (ZGA) of *Drosophila* embryogenesis^26–28^. To quantify its translation, we apply two parallel single-molecule imaging methods for mutual validation. The fraction of *hb* mRNA engaged in translation is spatially heterogeneous, exhibiting a plateau in the anterior region of embryos and a significant decline at the *hb* mRNA boundary, which differs from the *hb* mRNA spatial distribution and contradicts the conventional constant translation rate model. We further reveal that this heterogeneity of translation is *hb* mRNA concentration–dependent, with a ratio increasing and then saturating as mRNA concentration rises. Moreover, it represents a cross-scale framework that mRNA concentration–dependent translation contributes to tracking of protein abundance to declined mRNA levels, and sharp protein boundary, by relaxing the requirement of a short protein half-life value, especially under resource constraint during *Drosophila* embryogenesis.

## RESULTS

### Antibody-based detection validates the SunTag system for visualizing endogenous translation of *hb* before major ZGA

To link the contribution of the single-molecule translational kinetics to the dynamic protein patterning in embryos, we first quantify the translation of *hb* mRNAs across spatial scales in *Drosophila* embryos. The SunTag systems amplifies the translational signals by inserting a tandem array of 24 copies of GCN4 epitopes to the downstream of the *hb* start codon via clustered regularly interspaced short palindromic repeats (CRISPR) knock-in. To avoid effects from insufficient *in vivo* scFv-GFP detector supply (Fig. S1), we visualize SunTag amplification signals at the single-molecule level by freely-tunable and externally-provided anti-GCN4 antibodies. Accompanied with single-molecule fluorescence *in situ* hybridization (smFISH) probes targeting the *hb* mRNAs bearing *SunTag* sequence, we can detect both the translating and non-translating *hb* mRNA in single-molecule resolution (Fig. 1A, Methods).

**Figure 1.**
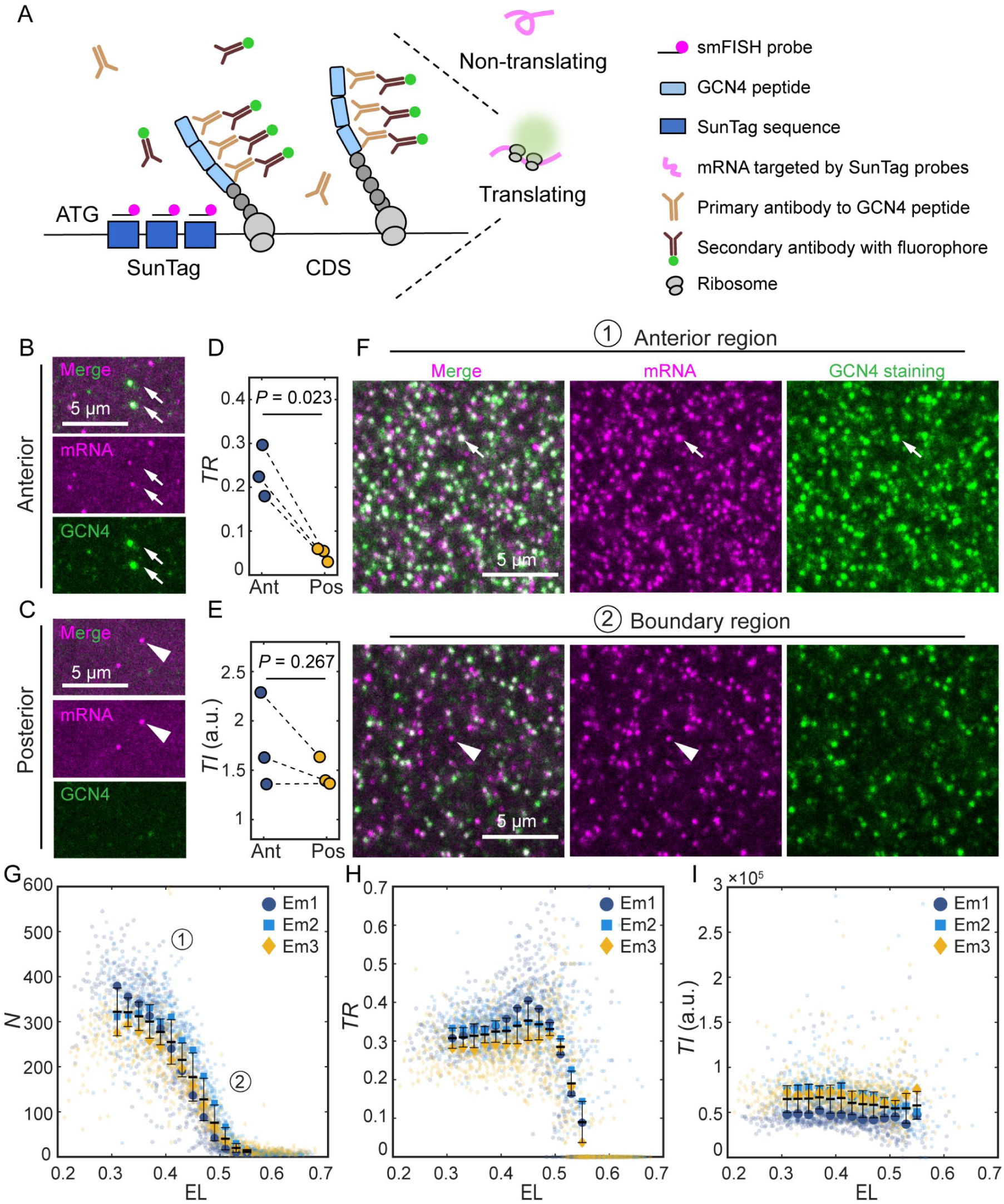
The anti-GCN4-based SunTag system reveals spatial heterogeneities of *hb* mRNA translation during embryogenesis. (A) Schematic of the SunTag system visualizing mRNA translation at single-molecule resolution. (B, C) Maximum intensity projection (MIP) of 21 successive confocal z-stack images of *hb-SunTag* embryos in nc 6–8, showing *hb* mRNA (magenta) and anti-GCN4 translational signal (green) in the (B) anterior and (C) posterior region. Subsets of *hb* mRNAs with (white arrows) and without (white triangles) co-localized translational signals are indicated. (D, E) Quantitative comparisons of the anterior and posterior regions along the anterior-posterior (AP) axis for (D) translational ratio (*TR*) and (E) translational intensity (*TI*) values, respectively (Welch’s *t*-test, sample size *n* = 3). Maternal *hb* transcripts from heterozygous *hb-SunTag* mothers are imaged in (B–E). (F) MIP of 3 successive confocal z-stack images of *hb-SunTag* embryos in nc 14, showing *hb* mRNA (magenta) and anti-GCN4 translational signal (green) in the anterior region (top panel) and boundary region (bottom panel) of the *hb* profile. (G, H, I) Spatial distributions of (G) *N*, mRNA concentration detected by *SunTag* smFISH, (H) *TR*, the ratio of mRNAs engaged in translation in nucleus volume unit, and (I) *TI*, total fluorescence intensity of translational signals over total translating mRNA concentration in nucleus volume unit along the AP axis, normalized to embryo length (EL) with 0 and 1 representing the anterior and posterior pole, respectively. Spatial heterogeneity of *TR* is assessed using a permutation-based *Kruskal–Wallis* (*KW*) test with Bonferroni correction (Methods), yielding adjusted *P*-values of approximately 3 × 10⁻⁴ across three biological replicates. Adjusted *P*-values of *post-hoc* pairwise comparisons are demonstrated in Fig. S3, Fig. S4, Fig. S5 (Methods). Points represent the mean values averaging from data in nucleus volume units (background points) in each bin from individual embryos (bin size: 0.02 EL). Sample sizes of *KW* test for each position bin in individual embryos are listed in Table. S1. Bold lines and error bars represent the means and standard deviations from points in each bin (bin size: 0.02 EL). Zygotic transcripts in heterozygous *hb-SunTag* (*w[*]; sp/Cyo; hb-SunTag/ TM6B, Tb[+]*) embryos are imaged in (F, G, H, I).

For validation, we quantify the posterior repression of maternal *hb* mRNA translation before ZGA (nuclear cycle, nc 6–8) reported in previous publications^26,27,29^. We define the translational ratio (*TR*) as the ratio between the number of the translating mRNAs (with both smFISH and translational signals) and total mRNAs (total smFISH signals) in a specific imaging volume consistent in anterior and posterior regions. *TR* of *hb* mRNAs in the anterior region ranges from 0.18 to 0.30 among replicates, while it drops significantly below 0.10 in the posterior region (Fig. 1D). Only a subset instead of all mRNAs are translating in a specific time point, in accordance with the stochastic nature of the translational reactions and the burst-like translation model^30^.

We next define the translation intensity (*TI*) as the total fluorescence intensity of the translational signals over total counts of mRNAs with translation signals in a specific volume. While *TR* reports the fraction of mRNAs engaged in translation, *TI* reflects the relative translational output of active mRNAs in a specific volume, i.e., translational efficiency. Based on the immunofluorescence (IMF) staining approach, *TI* values in the anterior region show moderate but not significant difference from those in the posterior region (Fig. 1E), suggesting that posterior repression primarily reduces the proportion of translating mRNAs rather than translational efficiency.

### Spatial heterogeneities of the single-molecule mRNA translational probability during major ZGA wave onset

We then apply this SunTag system to detect the *hb* translation during the onset of the major ZGA wave (∼ 6 min of nc 14) ^26,27,31^. Nuclei in syncytium embryos in this stage still lack cell membranes separation. Thus, the volume unit for the determination of *TR* and *TI* is determined by a Voronoi segmentation–like assignment of the single-molecule objects to their nearest nucleus. The *hb* mRNA (*hb* mRNA bearing *SunTag*) count in this nucleus volume unit is defined as *N*, analogous to the *hb-SunTag* mRNA concentration. We project *N*, *TR* and *TI* to the one-dimension AP axis for further quantification.

During this stage, *N* establishes a step-like patterns in the anterior region of the embryos, different from the evenly distribution before ZGA (Figure 1F, G). With smFISH probes targeting *SunTag* region, we employ *hb-SunTag* heterozygote (only one of two *hb* alleles bearing *SunTag* insertion, called *hb-SunTag* fly line hereafter) to decrease the density of detected mRNAs for the convenience of image processing (Methods). The mRNA concentration after scaling to *N*_scaled-s_ (Fig. S2, Methods) coincides with the published values detected in wildtype^32^.

Besides mRNA pattern, *hb* translation ratio does not keep constant as usually assumed during the onset of the major wave of ZGA, while it is also distinct from posterior repression manner observed in maternal *hb* mRNAs (Fig. 1F, H, S3–5). Specifically, the *TR* value is 0.09 ± 0.05 at around 0.55 ± 0.01 EL near the boundary region of mRNA profiles. This ratio increases gradually and approximately reaches a constant towards anterior region, with the representative value of 0.30 ± 0.02 at 0.31 ± 0.01 EL averaged among three replicates (Fig. 1 H). We further demonstrate that both the values and spatial patterns of *TR* along the AP axis are maintained the same when the dilution ratio of antibody detectors is varying from 1:1000 to 1:100 (Fig. S6), indicating the independency of the *TR* heterogeneity under ordinary antibody concentration. Based on our antibody staining approach, *TI* is unlikely to be spatially heterogeneous along the AP axis (Fig. 1I).

### The genetic manipulation–free measurement of *hb* mRNA end-to-end distance supports spatial heterogeneity of translation

To rule out potential artifacts from genetic perturbations and further validate the detection of translation heterogeneity, we develop a genetic-manipulation–free approach. Previous studies have demonstrated that the binding of ribosomes to mRNA contributes to mRNA structural extension, leading to a positive correlation between translation efficiency and mRNA end-to-end distance^33–36^. We label the 5′ and 3′ ends of *hb* mRNA directly in embryos using two sets of single molecule inexpensive FISH (smiFISH) probes with two distinct fluorophores, and employ stimulated emission depletion (STED) microscopy to measure mRNA end-to-end distances.

To rigorously validate that our STED-based measurements could resolve conformational differences of mRNAs at various translation states, we first establish the spatial resolution of our STED imaging system under experimental conditions. Using dual-color probe sets targeting the same region of the *hb* transcript (co-localization control, Fig. 2A up panel), we measure a median distance of 34 nm (Fig. 2B), indicating that our STED setup can reliably distinguish end-to-end distances exceeding this threshold. We further assess the localization precision of this imaging setting by fitting the point-spread function to representative single-molecule objects in each channel. The calculated localization precision approaches the default spatial resolution of STED microscope (AF594 channel: X direction: 27 ± 4 nm, Y direction: 37 ± 10 nm; ATTO647N channel: X direction: 29 ± 3 nm, Y direction: 28 ± 9 nm).

**Figure 2.**
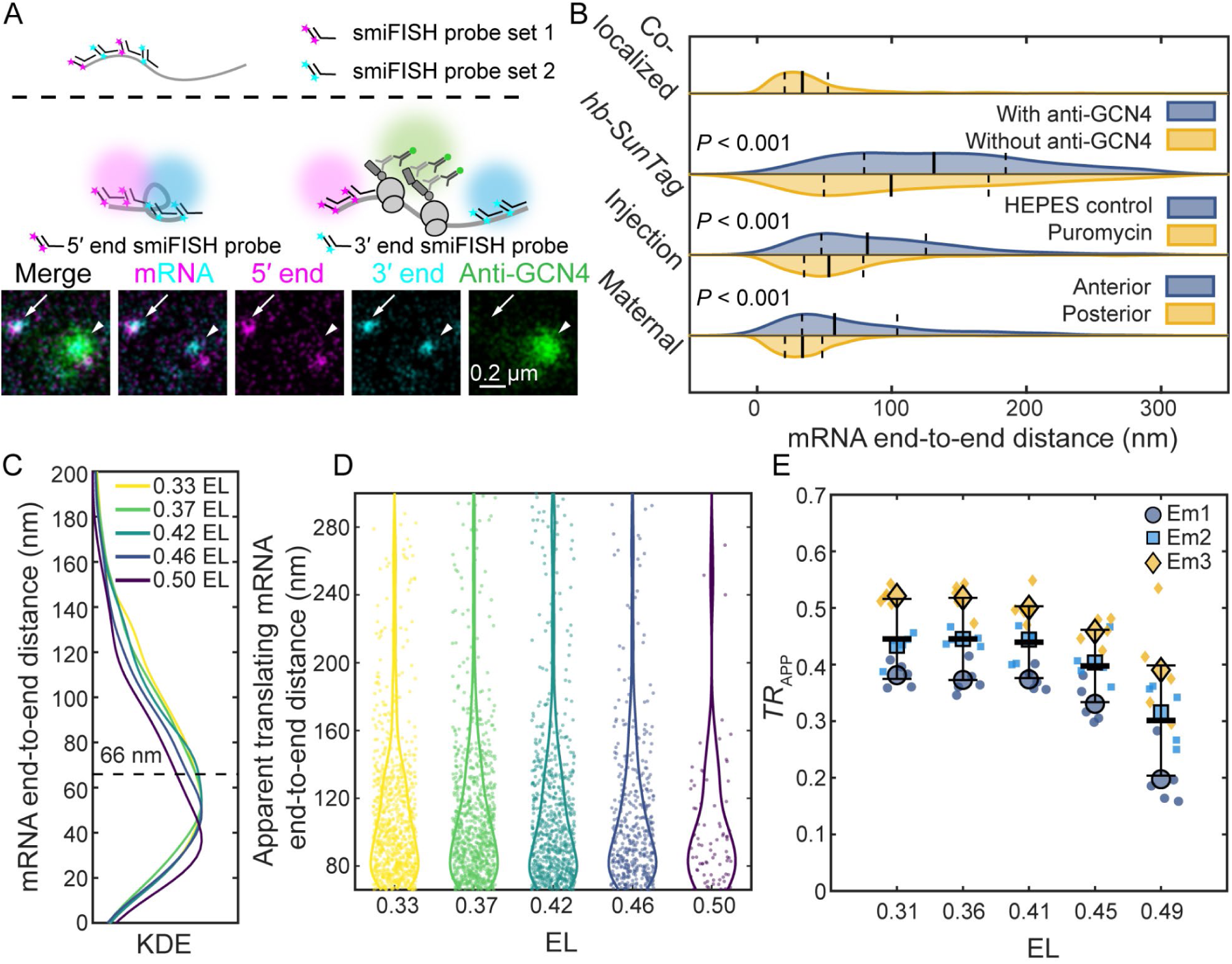
The measurement of the mRNA end-to-end distance also reveals spatial heterogeneity of *hb* mRNA translation during embryogenesis. (A) Upper panel: Schematic of the measurement of dual-label co-localization control experiment. Two probe sets target a same region on hb mRNA. Middle panel: Schematic of the measurement of end-to-end distance of zygotic *hb* transcripts in *hb-SunTag* (*w[*]; sp/Cyo; hb-SunTag/ TM6B, Tb[+]*) heterozygous embryos. The 5′ and 3′ ends of *hb* mRNAs are targeted with a smiFISH probe set, respectively. Each single smiFISH probe contains an “L”-shaped primary probe targeting mRNA sequence, and a fluorophore-labeled secondary probe recognizing the free arm of the primary probe (supplement text). Lower panel: representative images of zygotic *hb* mRNA end-to-end measurement in *hb-SunTag* heterozygous embryos. White triangles and arrow represent the mRNA with and without anti-GCN4 translational signal co-localization. (B) KDEs corresponding to dual-label colocalization experiment, end-to-end distance measurement of *hb-SunTag* mRNAs, puromycin injection experiment, and end-to-end distance measurement of hb mRNA in in *w^1118^* embryos in nc 6–8. Welch’s *t*-test. Sample size for each experiment *n* = 1041, 1034, 289, 397, 689, 513, 328. (C) Truncated KDEs (whole KDEs: Fig. S7 C, right panel) of *hb* mRNA end-to-end distance from 5 anterior-posterior (AP) positions in a representative embryo in nc 14. The AP axis is normalized to embryo length (EL) with 0 and 1 representing the anterior and posterior pole, respectively. Dashed line annotates the determined distance threshold (66 nm). (D) Violin plots for translating mRNA end-to-end distances for *hb* mRNAs (background points) from 5 AP positions (*n* = 853, 870, 736, 415, 77. The adjusted *P*-value of the permutation-based *KW* test with Bonferroni correction working as a statistical test for the spatial heterogeneity of end-to-end distance of translating mRNAs: 0.800.) in the corresponding embryo in (C). (E) The spatial heterogeneity of distance inferred *TR*_App_ determined by the mRNA end-to-end distance in a specific volume along the AP axis in 3 embryos. Points represent the mean values averaging from imaging areas (background points) at each position in individual embryos. *n* = 5 of permutation-based *KW* test with Bonferroni correction works as a statistical test for the spatial heterogeneity of *TR* in individual embryos. Adjusted *P*-values for 3 replicates: 0.001, 0.008 and 0.018. Bold lines and error bars represent the means and standard deviations from points in each AP position. (B) Maternal and (A, C, D, E) zygotic transcripts in *w^1118^* embryos are imaged.

We next assess whether translation status correlates with mRNA extension using the *hb-SunTag* reporter system, which allows simultaneous detection of mRNA ends and nascent peptide signals (Fig. 2A, bottom panel). Consistent with prior observations that ribosome occupancy contributes to mRNA structural extension^33–36^, *hb* mRNAs co-localizing with anti-GCN4 signals, which is an indicative of active translation, exhibit significantly greater end-to-end distances (median: 131 nm) compared to those lacking nascent peptide signals (median: 100 nm) (Fig. 2B). Although the two populations exhibit some overlap, the distribution of mRNAs without anti-GCN4 signal co-localization is markedly enriched for shorter end-to-end distances, supporting the positive correlation between translation and mRNA extension.

To further confirm that the observed compaction is a direct consequence of translational activity rather than primarily by RNA binding protein (RBP) binding or other structural constraints, we treat wild-type *w^1118^* embryos in nc 14 with puromycin, which induces premature chain termination and ribosomal dissociation. Puromycin treatment results in a pronounced shift toward shorter mRNA end-to-end distances (median: 53 nm) compared to the control group with HEPES buffer injection (median: 82 nm) (Fig. 2B), arguing against the possibility that the extended conformation observed in the anterior region of nc 14 *w^1118^* embryos is primarily driven by RBPs without effect on translation, as such binding would not be expected to produce puromycin-sensitive structural changes. To further support that *hb* mRNA structure extension is related with translation in our case, we exploited the naturally occurring translationally repressed maternal *hb* mRNA pool in the posterior region believed to be mediated by RBP binding. The mRNA end-to-end distance of *hb* mRNAs in the posterior region of embryos in nc 6–8 also exhibits an enrichment in the shorter distance range (median: 34 nm), with Kernel density estimation (KDE) of posterior-region end-to-end distances reveals a sharp decline below 100 nm, in contrast to the more gradual distribution in the anterior region (median: 58 nm) (Fig. 2B).

For simplification in calculating the translation ratio as in the SunTag measurement, we derive a threshold for operationally distinguishing translating from non-translating mRNAs in nc 14 embryos. We identify the threshold distance by locating the maximum non-zero peak of the second derivative of the posterior KDE, which marks the steepest point of decay to represent the most prominent difference between the posterior (mostly non-translating) and anterior (translating mixed with non-translating) profiles. This analysis yields a threshold of 66 nm, averaged across three biological replicates (Fig. S7A, B). Notably, the co-localization precision control (dual probes on the same region, Fig. 2B) produced a distance distribution closely resembling that of posterior maternal mRNAs (median: 34 nm, Fig. 2B), confirming that the 66 nm threshold lies well above the technical resolution limit and thus reflects bona fide biological compaction rather than imaging artifacts.

Applying this 66 nm threshold to nc 14 embryos, we calculate the distance-inferred apparent translation ratio (*TR_App_*) along the AP axis. The KDE profiles of end-to-end distances during the major ZGA wave exhibited clear spatial variation (Fig. 2C, Fig. S7C). Consistent with our SunTag-based measurements (Fig. 1H), *TR*_App_ remained relatively stable across the anterior *hb* expression domain and decreased significantly near the posterior boundary (Fig. 2E). Quantitatively, *TR*_App_ was 0.45 ± 0.07 at 0.31 ± 0.02 EL, and 0.30 ± 0.10 at 0.49 ± 0.01 EL averaged across replicates (Fig. 2E; adjusted *P*-values from *post-hoc* pairwise comparisons are shown in Fig. S8; see Methods).

Importantly, when we restrict our analysis to mRNAs with end-to-end distances exceeding 66 nm, corresponding to those apparently classified as translating, we find no significant variation in their distances along AP axis (Fig. 2D; Fig. S7D, E; see Methods). This observation, together with *TI* measurements from the SunTag system, further supports that translational efficiency can be uniformly distributed along the AP axis, whereas the fraction of actively translating mRNAs (*TR*) displays spatial heterogeneity.

### *TR* heterogeneity exhibits mRNA concentration–dependence

A straightforward explanation for the spatially heterogeneous *TR* of *hb* along the AP axis would be a positional gradient cue, such as a repressive signal emanating from the posterior side of the embryo. However, we find that the local heterogeneity in both *TR* and *N* can be unified into a single, position-independent relationship. Strikingly, irrespective of the AP position, *TR* exhibits a clear dependence on *N*, suggesting that local mRNA concentration, not the AP coordinate, encodes spatial heterogeneity in translation of *hb*.

We firstly observe a non-linear relationship between *TR* and *N* values (Fig. 3A). Binning along mRNA concentration *N* (bin size: 20), *TR* increases from 0.24 ± 0.07 at the *N* value of 13 ± 10 and start to saturate as *N* equaling to 73 ± 10, and end at *N* equaling to 333 ± 10, with corresponding *TR* value of 0.43 ± 0.04 (Fig. 3A). To quantify this relationship, we develop an mRNA concentration– dependent model relating *TR* increasing to *N* changes (eq1),

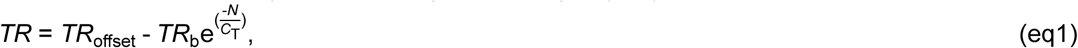

where *TR*_offset_ represents the global *TR* offset value, *TR*_b_ determines the increasing amount of *TR*, and *C*_T_ determines the concentration constant reflecting the curve steepness of *TR* reaching saturation. This model properly fits the increasing-saturation change of *TR* as *N* elevates (Fig. 3A, Methods, Table S2).

**Figure 3.**
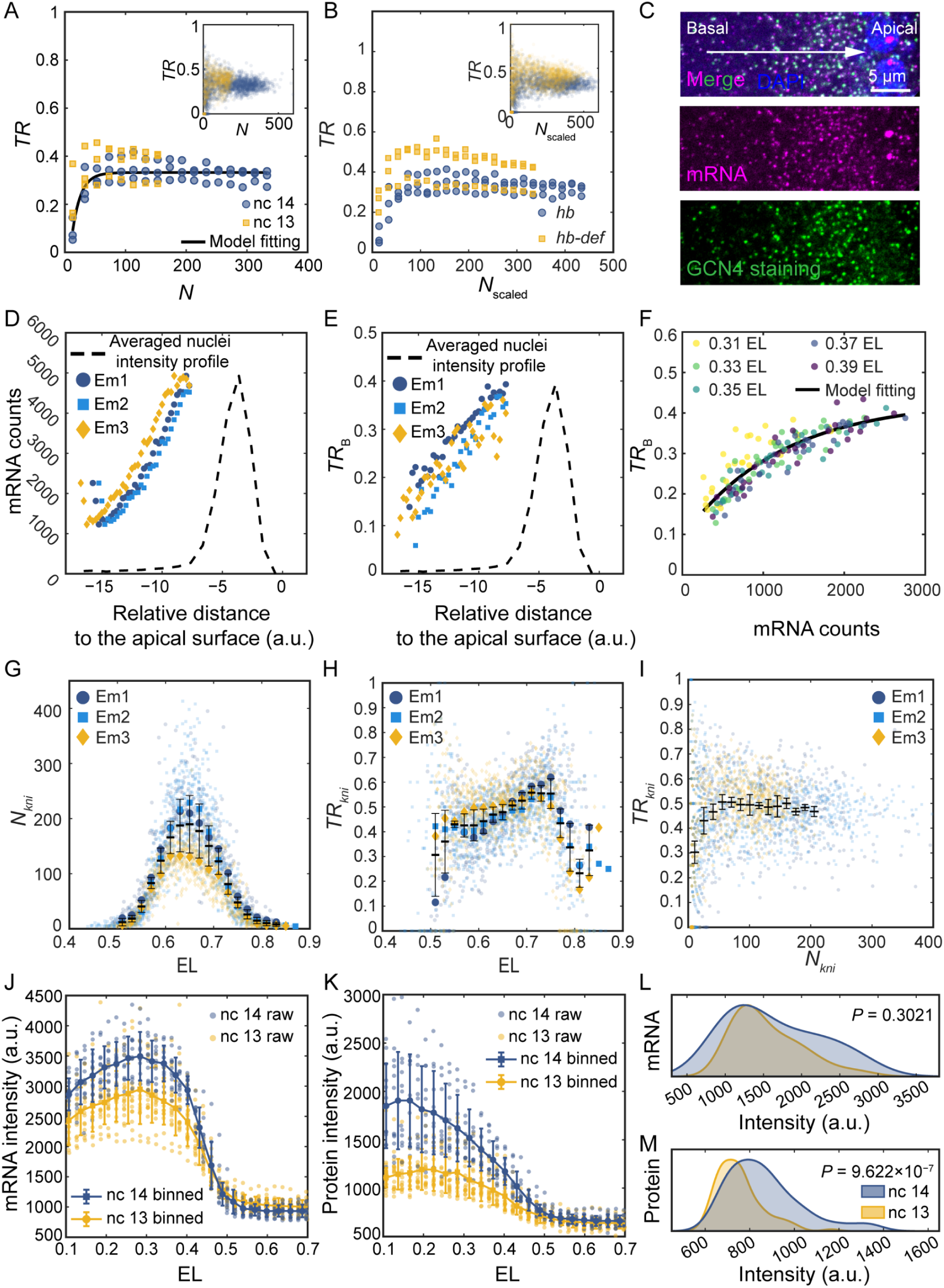
The dependence of the ratio of *hb* mRNAs engaged in translation on *hb* mRNA concentration. (A) The *N*-*TR* (mRNA concentration-ratio of mRNA engaged in translation) relationship in nc 13 (sample size *n* = 3) from the SunTag system agrees with the prediction from mRNA concentration–dependent model fitting to the *N*-*TR* data in nc 14 (*n* = 3). Points represent the mean values averaging from the nucleus volume units (data points in inset) in each bin from individual embryos (bin size: 20). (B) The *N*_scaled_-*TR* relationship between *hb-SunTag* and *hb-def* fly lines. The *N* values of *hb-SunTag* are scaled to total *hb* transcript concentration (*N*_scaled_) in embryo for the comparison with *hb-def* of which *N* equals to total *hb* transcript concentration (*N* = *N*_scaled_). Points represent the mean values averaging from the values in nucleus volume units (data points in inset) in each bin from individual embryos (bin size in *N*_scaled_ value: 20). A scaling factor of 1.31 is applied to *N* of heterozygote *hb-SunTag* fly line to account for the contribution of the untagged balancer *hb* allele in this fly line. This factor was determined by calculating the ratio of total hb mRNA counts to *SunTag*-bearing *hb* mRNA counts from representative images of each embryo (1.32, 1.33, 1.28). (C) Maximum intensity projection (MIP) of 3 successive confocal z-stack images of a representative *hb-SunTag* embryo (nc 14) showing nuclei (blue), *hb* mRNAs (magenta) and translational signals (green) in the basal region. (D) Total mRNA counts and (E) *TR*_B_ increase when approaching the nuclei (step size: 0.25 μm) in three replicates. Nuclei intensity from 3 replicates are averaged and normalized based on the exact mRNA counts or *TR*_B_ values for illustration. (F) The mRNA concentration–dependent model can also account for the relationship between *TR*_B_ and mRNA count in the basal region at multiple AP positions (represented by colored points, embryo number *n* = 3). (G, H) Spatial distributions of (G) *N_kni_*, *kni* mRNA concentration detected by *SunTag* smFISH, (H) *TR_kni_*, the ratio of *kni* mRNAs engaged in translation in nucleus volume unit. (I) The non-linear relationship between *N_kni_* and *TR_kni_*. (J, K) The (J) *hb* mRNAs and (K) Hb proteins are simultaneously quantified by combining smFISH with probes targeting the *hb* coding sequence and anti-Hb IF staining, respectively, in individual embryos in nc 13 (*n* = 17) and early nc 14 (*n* = 12). The anterior-posterior (AP) axis is normalized to embryo length (EL) with 0 and 1 representing the anterior and posterior pole, respectively. Background points: averaged fluorescence intensity of the pixels at each AP bin (bin size: 0.02 EL) in individual embryos. Points and error bars: mean and standard deviation of the background points among embryos (bin size: 0.03 EL). (L, M) Comparing the fluorescence intensity of (L) *hb* mRNA and (M) Hb protein in 0.42–0.53 EL between nc 13 (*n* = 102) and early nc 14 (*n* = 72), Welch’s *t*-test. Zygotic (nc 14) and dominantly zygotic (nc 13) transcripts in *w^1118^* embryos are imaged in (J–M) (A, D–F) Zygotic transcripts in heterozygous *hb-SunTag* (*w[*]; sp/Cyo; hb-SunTag/ TM6B, Tb[+]*) embryos are imaged. (B) Zygotic transcripts in heterozygous *hb-SunTag* (*w[*]; sp/Cyo; hb-SunTag/ TM6B, Tb[+]*) and *hb-def* (*w[*]; sp/Cyo; hb-Df(3R)BSC197/ TM6B, Tb[+]*) embryos are imaged. (G–I) Zygotic transcripts in heterozygous *kni-SunTag* (*w[*]; sp/Cyo; kni-SunTag/ TM3, sb*) embryos are imaged.

To decouple the effect of spatial position and mRNA concentration, we first test how changes in *N* values influence *TR* by perturbing *hb* mRNA concentration. Taking advantage of the naturally lower *hb* mRNA concentration at earlier time point nc 13 (∼50% of that at nc 14), we quantify the *N*-*TR* relationship of *hb* in the *hb-SunTag* fly line, which maintains a consistent genetic background. In agreement with our model’s prediction, this *N*-*TR* relationship at nc 13 retracts towards a lower mRNA concentration region (Fig. 3A).

Moreover, while keeping the developmental time point constant at nc 14, we quantify *TR* in a single-*hb-*allele deficient mutant (*hb-def*), which has ∼76% of the mRNA concentration in the *hb-SunTag* fly line (see Methods for details). We still observe a similar *N*-*TR* increasing-saturation relationship in the *hb-def*, again shifting towards the lower mRNA concentration region (Fig. 3B). This result demonstrates that the mRNA concentration–dependent relationship between *TR* and mRNA concentration is preserved in this mutant, albeit with a moderate shift that may arise from differences in genetic background affecting baseline levels of the translational regulators and machineries between two fly lines (*hb-SunTag* and *hb-def*).

We further decouple the *N*-*TR* relationship from spatial context by examining the basal region of the embryo cortex. Consistent with the previous study^32^, we observe a *hb* mRNA density gradient in the basal region of cortex, with higher level beneath the nuclei and decreasing towards yolk (*z* step size: 0.25 μm, Fig. 3C, D). Remarkably, rather than remaining constant, the translational ratio in each basal step, *TR*_B_, also exhibits heterogeneity along the basal region (Fig. 3E). Zooming into different AP positions, *TR*_B_ also follows a *N-TR*_B_ relationship. Though with distinct parameter values, our mRNA concentration–dependent model still accurately described the relationship between mRNA counts and *TR*_B_ in a spatial context different from AP axis profiles (Fig. 3F, Table S2). This result provides additional evidence of local mRNA concentration, rather than spatial coordinates that predicts the translational ratio.

Most importantly, the difference of *TR* of *kni* provides particularly compelling evidence for mRNA concentration–dependent translation. Unlike *hb*, whose *TR* declines only toward the posterior as its mRNA concentration decreases (Fig. 1G, H), *kni TR* decreases on both sides of its posterior mRNA peak (Fig. 3G, H). This pattern rules out a single unified spatial gradient cue: if a posterior-derived repressive signal governed translation, *TR* should decline specifically toward the posterior in both cases, not on both sides of the *kni* peak. Explaining this under a purely spatial regulatory model would require two opposing cues, one anterior and one posterior, each suppressing translation in opposite directions. However, given the narrow width of the *kni* peak, these two cues might be sophisticatedly separated to prevent possible overlap inside the *kni* mRNA peak to maintain the observed high *TR* at approximately the peak. This model requires more complex processes and patterning, and thus seems less likely to be the primary regulation. In contrast, both *hb* and *kni* collapse onto essentially the same *TR*-*N* relationship despite their distinct spatial patterns (Fig. 3I), representing a simple unified model. These results collectively supporting local mRNA concentration as the potential determinant of translational regulation.

To further test whether mRNA rather than protein drives *TR*, we compared mRNA and protein profiles between nc 13 and nc 14. (Fig. 3J, K). We conduct smFISH for *hb* mRNA and IMF for Hb protein on *w^11^*^18^ embryos at nc 13 and nc 14 in a single batch, and quantitatively analyze the cytoplasmic mRNA and protein level in two developmental stages. The mRNA level along the AP axis should also exhibit overlap between nc 13 and nc 14 before their corresponding *TR* reaching plateaus value in *N-TR* curve, if *TR* changed is compatible with *N*-*TR* relationship. Indeed, the *hb* mRNA expression levels in nc 13 and nc 14 overlap in 0.42–0.53 EL (Fig. 3J, L). Comparing with the cytoplasmic Hb protein level in nc 13 and nc14 (Methods), the protein levels between these two stages are divergent in the same AP axis region as for mRNA profile (Fig. 3K, M). This result supports the decoupling of *N-TR* relationship from the AP axis context.

Finally, we assess the spatial distribution of one of a ribosomal protein tagging with GFP, RpL10A-GFP. RpL10A-GFP is evenly distributed along AP axis and in the basal region under our ongoing imaging and analysis procedure, reducing the possibility of explaining the *TR* pattern by spatial distribution of ribosomes (Fig. S9). But we emphasize that the absence of spatial heterogeneity in RpL10A-GFP is not equal to the evenly distribution of ribosomes. The spatial heterogeneity in two other ribosomal proteins reported in previous works indicates the possibility of specific enrichment for free ribosomal proteins^22^.

Collectively, our results highlight mRNA concentration as a highly possible predictor of *TR* heterogeneity. The mRNA concentration–dependent regulation of *TR* is conserved across different gap genes, developmental stages, and embryo axes. Although model parameter values vary, the core functional form of the *TR-N* relationship consistently explains the data.

### mRNA concentration–dependent model contributes to Hb protein profile establishment across spatial scales

Hb is a key transcription factor in the segmentation gene network, which establishes highly dynamic protein pattern with a sharp boundary under a tight resource budget^6,37,38^.

To assess the potential contribution of mRNA concentration–dependent translation to Hb patterning, we perform computational simulations comparing this mechanism against the conventional model assuming a constant translation rate (briefed as constant model hereafter). Using published temporal profiles of *hb* mRNA and protein^6^, we fit both mathematical frameworks to the experimental protein data with mRNA data input, and evaluate each model’s ability to recapitulate the observed Hb dynamics. The mRNA and protein levels are firstly projected to the AP axis, with *Pro*_i_ and *m*_i_ representing the protein and mRNA levels at AP position *i* (representing 0.01 EL bin), while *D* (0.01 EL^2^ / min) and *λ* (min^−1^) representing the protein diffusion constant and degradation rate (eq2 and eq3), respectively. In the constant translation model, *k* (min^−1^) represents constant translational rate (eq2)^39,40^. In the mRNA concentration–dependent model, *TR* implicitly multiplies a constant *TI* as the translation rate with *k*_offset_ (min^−1^) and *k*_b_ (min^−1^) in eq3 related to *TR*_offset_ and *TR*_b_ in eq1.

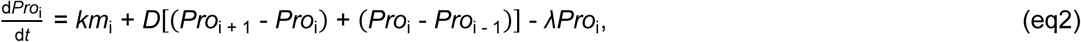

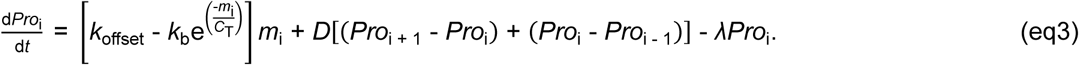

Among all tested protein half-life values, the mRNA concentration–dependent translation model (Eq. 3) consistently yields the best fit to both the experimentally observed Hb boundary sharpness and boundary positions (Fig. 4A, Methods, Table S3). In contrast, the constant translation model produces only a modest increase in simulated boundary sharpness when protein degradation rates are elevated. This indicates that, within the conventional constant model, higher degradation rates can partially preserve boundary sharpness, but at the expense of increased protein turnover and thus elevated resource expenditure. The superior performance of the mRNA concentration–dependent model highlights the global contribution of this translation mode to establishing protein boundary sharpness under a resource-constrained system.

**Figure 4.**
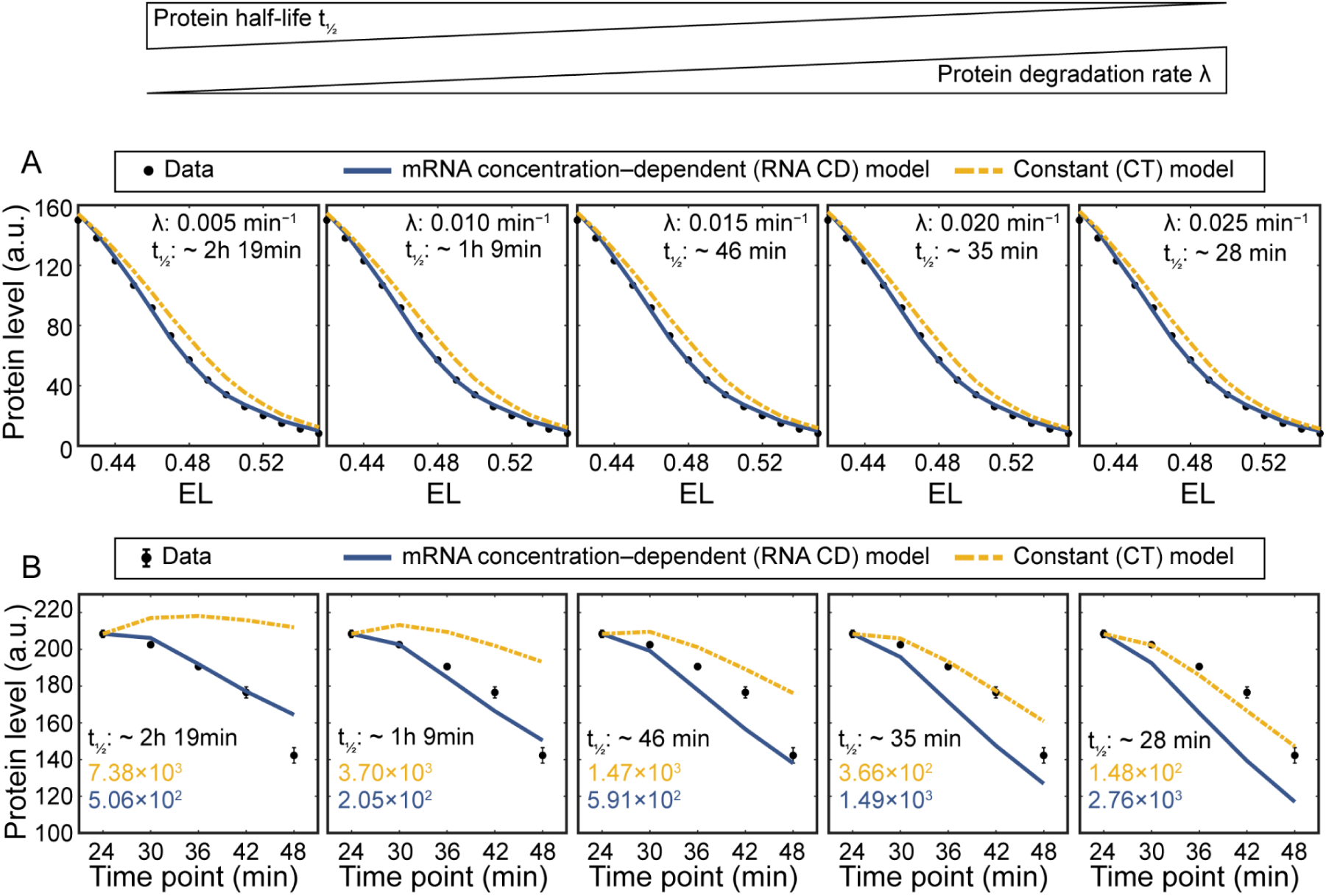
The mRNA concentration–dependent translation model contributes to rapid establishment of the Hb protein profile with sharp boundaries at low resource cost. With protein half-life ranging from ∼ 2h 19min to ∼ 28 min, (A) the mRNA concentration–dependent model outperforms the constant model in fitting the experimental data of the Hb protein profile along the AP axis at a subsequent time point (∼ 12 min into nc 14) of the onset of nc 14. (B) Comparison of the residual sum of squares (RSS) of sequential between experimental data and model predictions of the Hb protein level averaging from 0.27–0.33 EL. The advantage of the mRNA concentration– dependent model becomes more pronounced with increasing protein half-life. Points and error bars represent the means and standard deviations. The experimental data of *hb* mRNA and Hb protein is from reference 6.

Besides the boundary sharpness, the mRNA concentration–dependent translation enables faster response of protein to declining mRNA levels during development without the extra cost of the short protein half-life required by the constant model (Fig. 4B). The proper decline of developmental pattern is of equally important to its build-up. Experimentally, reductions in *hb* mRNA concentration at 0.27–0.33 EL during later time points (∼ 24–48 min) at nc 14 give rise to a two-stripe mRNA pattern and a concomitant decrease in protein abundance^6^. The proper Hb protein level control in response to mRNA concentration decreasing in this region is crucial for the following developmental procedures. Inadequate reduction of Hb protein in this AP axis region leads to distortion of downstream developmental patterns driven by extra Hb, represented by expanded *even-skipped* (*eve*) mRNA stripe 1 located at ∼ 0.30 EL^41^. Comparing two models, the sequential protein level changes simulated from mRNA concentration–dependent model in 0.27–0.33 EL in later time points of nc 14 responds to the *hb* mRNA abundance reduction faster in all tested protein half-lives (Fig. 4B, Methods). By evaluating the residual sum of squares (RSS) between experimental and simulated data, we find that mRNA concentration–dependent model outperforms the constant-translation model in explaining the observed protein dynamics under longer protein half-life conditions. A longer half-life enhances the resource efficiency of protein expression, as each synthesized molecule remains functional over an extended period. Thus, the mRNA concentration–dependent translation correspondingly reduces the protein synthesis as the mRNA level decreasing. As protein half-life becomes shorter, the constant-translation model appears to fit well with the experimental data at the cost of high resource expenditure. These results further support the conclusion that mRNA concentration–dependent translation alleviates the resource efficiency constraint, thereby enabling the co-existence of dynamic and sharp protein pattern in *Drosophila* embryos.

In summary, the mRNA concentration–dependent translational regulation relaxes the inherent constraint of the paradox between protein pattern pace and reducing resource burden for embryo development (Fig. 5). In the conventional constant translation rate framework, sharp boundary and faster response of protein to the mRNA abundance change are correlated to shorter protein half-life, which increases the resource expenditure during protein turnover. However, the mRNA concentration–dependent translation effectively compensates parts of the requirement for high protein degradation rate. This regulation at single-molecule level ultimately facilitates the responsiveness and boundary sharpness of protein patterns at the whole-embryo spatial scale, comparing with the constant translation (Fig. 5).

**Figure 5.**
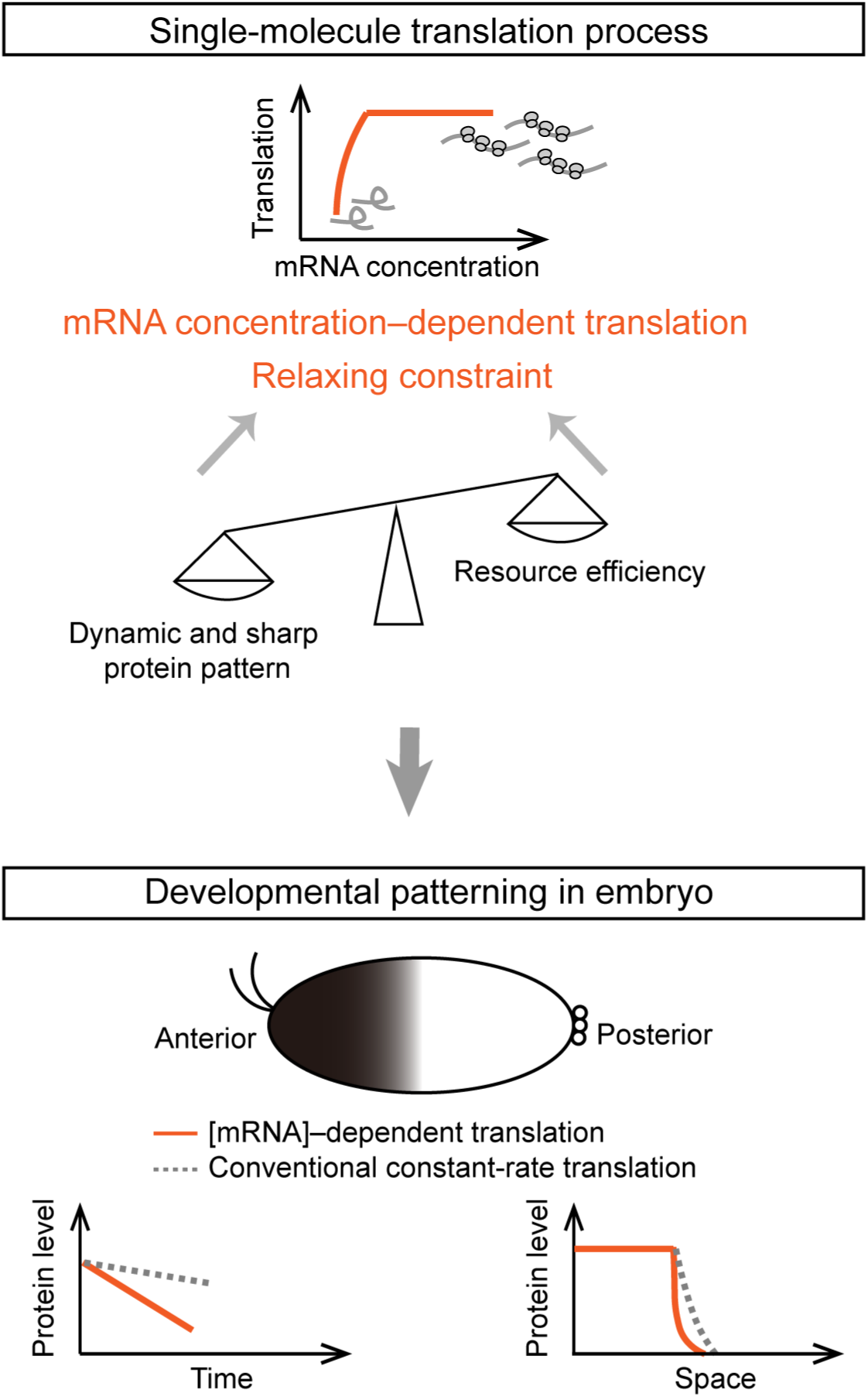
Schematic illustrating how mRNA concentration–dependent translation relaxes the resource constraint to enable fast protein response and sharp Hb boundary formation.

## DISCUSSION

In this work, we reveal how an mRNA concentration–dependent translational relationship can relax the paradox between the need for fast sharp protein patterning and the underlying resource constraint during *Drosophila* embryogenesis. Hb is an essential transcription factors for segmentation, and its protein pattern must be established rapidly and precisely within a limiting developmental time window. Our results demonstrate that *hb* mRNA is not translated under a constant rate across the embryo. Instead, its translational probability changes with local mRNA concentration, preserving the responsiveness of protein production while avoiding unnecessary synthesis. This represents an economical use of limited resources, as protein production is prioritized where mRNA is scarce and most rate-limiting for patterning. Compared with spatially encoded regulatory cues, which would require distinct anterior and posterior signals acting on different targets, a unified mRNA concentration–dependent rule offers a simpler solution: the same regulatory logic operates across genes and positions without invoking multiple position-specific mechanisms. This strategy thus provides a potential design principle by which developmental systems achieve fast and sharp protein patterning under biosynthetic resource constraint and stringent temporal limits.

This potential regulatory strategy carries important biological implications that production of RNA and protein imposes resource costs^9^, and such costs can shape the evolution of gene-expression strategies^1,42^. For instance, a trade-off between cell-to-cell gene expression precision and transcription cost has been observed across multiple organisms, where combinations of extreme high precision and large transcription cost are disfavored because they improve precision at the expense of excessive RNA production^1^. Recent work further suggests that the subcellular localization of mRNA and protein species to whether soma or dendrites adheres to principle of energy minimization^43^. In this context, our work offers a distinct prospective that embryos could modulate at the translation level according to the local mRNA level, a strategy that may be particularly useful in resource-constrained systems.

Multiple non-mutually exclusive mechanisms could potentially explain this relationship between translation and mRNA concentration. RBP-mediated regulation is a plausible mechanism. Early *Drosophila* embryos contain a large and developmentally dynamic RNA-bound proteome, and RBP interactome studies show extensive remodeling during MZT, suggesting that RBPs participate broadly in regulating protein production at this stage. Classical examples including posterior repression of maternal *hb* by RBPs and Me31B-mediated repression of maternal mRNAs demonstrate that *Drosophila* embryos possess RBP–RNA interactions capable of controlling protein expression with high specificity. Importantly, concentration changes in components of RBP– RNA complexes can give rise to emergent behaviors such as phase separation or condensate formation. For instance, FXR1 condensation enhances translation of stored mRNAs during spermiogenesis, and a recent synthetic system in *E. coli* showed that sequestering mRNAs into ribonucleoprotein condensates enhances their translation. These findings resonate with our observation that increased mRNA abundance correlates with increased translational activity, supporting a model in which higher local mRNA concentrations promote the formation of translationally favorable RBP–RNA assemblies. Other mechanisms may also contribute. Variations in *hb* mRNA abundance could influence access to limiting translational machinery, as previous experimental and theoretical work has shown that competition for ribosomes or tRNAs can link transcript concentration to protein production. Under this scenario, increased local *hb* mRNA concentration would enhance the effective availability of translational machinery to each transcript. Additionally, cis-regulatory elements—such as poly(A) tail length, which positively correlates with translational efficiency during early Drosophila embryogenesis, or RNA secondary structure and upstream open reading frames, which tend to interfere with translation—could modulate the relationship. Notably, trans- and cis-regulatory mechanisms often act concomitantly, and their interplay may further shape the concentration-dependent behavior we observe.

Future studies should explore the detailed underlying mechanism of the observed mRNA-concentration–dependent translation employed by *Drosophila* embryos. Systematically, multiplexed perturbations of components in translational machinery will be important for identifying candidate regulators for mRNA concentration–dependent translation. More directly, a profiling of the RBP interactome specific to *hb* mRNA could be implemented to identify proteins associated with *hb* mRNA, and to determine whether these associations vary with transcript abundance, time, and space. Correspondingly, cis-regulatory analysis of *hb* mRNA should determine whether the untranslated region (UTR) and coding sequence, or their combinations encode the nonlinear translational response. Combining the screenings and perturbations at molecular scale with the inspection of *Drosophila* embryo phenotypes, including Hb boundary sharpness, boundary dynamics, and segmentation precision and reproducibility, can bridge between molecule mechanisms to developmental contributions.

To further test the universality of the discovered translational regulation mechanism, several future directions are worth following from this work. Though it is technologically more difficult than reducing *hb* mRNA concentration in embryos, we consider to increase mRNA abundance, in either embryos or embryo extractions, to extend this mRNA concentration–dependent relationship to a broader concentration range. It will also be important to test whether similar mRNA concentration–dependent translational relationships occur in other developmental regulators, firstly in *Drosophila* embryos, for example, the gap gens other than *hb* and *kni* measured in this study, then to embryos in other species. If so, this mechanism may represent a general strategy by which embryos balance protein-pattern precision, developmental speed, and biosynthetic economy.

In conclusion, translation emerges as a critical and information-rich node in development. Although translational regulation is well-established as a key regulator at maternal level in *Drosophila* embryos, its functional significance and regulatory logic during the zygotic stages still remain incompletely understood. Notably, previous studies in *Drosophila* embryogenesis have predominantly focused on transcriptional networks and morphogen gradients^38,44,45^. Our discovery of mRNA concentration–dependent translation now adds a pivotal new regulatory block, demonstrating the coordination of translation by mRNA level across cells. This finding resonates the recently proposed “Cellular Dogma” which calls for conceptual framework beyond information transmission at the molecular level in classical central dogma, emphasizing instead the orchestration of information within and between cells in complex systems^46^.

## METHODS

### Cloning and transgene

For CRISPR knock-in of *SunTag* sequence to downstream of the *hb* start codon, sgRNA target sequence were designed using the FLYCRISPR Target Finder (http://targetfinder.flycrispr.neuro.brown.edu/). The pCFD1-dU6:1 plasmid (Addgene plasmid # 49408), a gift from Dr. Simon Bullock, was linearized by BbsI and ligate with the annealed oligonucleotides containing sgRNA sequences GGGCGATTGGCGCGGACTGC|TGG for *hb* and GGATTAAACCCTTACTAACG|CGG for *kni*. Donor plasmids for *hb-SunTag* and *kni-SunTag* CRISPR knock-in fly lines containing the SunTag epitope sequence and two 1500 bp homologous arms from *hb* and *kni* respectively. Two homologous arms flanking the ATG of targeted genes were PCR amplified from *w^11^*^18^ genomic DNA. The SunTag epitope sequence was PCR amplified from pcDNA4TO-24xGCN4_v4-sfGFP (Addgene plasmid # 61058), a gift from Dr. Ron Vale. The PCR amplification products, including two homologous arms from targeted genes and *SunTag* sequence, were fused by fusion PCR, and ligated into a linear T vector (pEASY-Blunt Simple Cloning Kit, TransGen Biotech) to construct the *hb-SunTag* donor plasmid, and ligated into a pUc19 vector to construct the *kni-SunTag* donor plasmid. The PAM site on *hb-SunTag* donor plasmid was mutated from CCA (TGG) to CAA (TTG). The PAM site on *kni-SunTag* donor plasmid was mutated from CCG (CGG) to ACG (CGT). The sgRNA plasmid and donor plasmid were co-injected into *y sc v; nos-Cas9; attP40* (TH00788.N) embryos by Tsinghua Fly Center in Tsinghua University. The expression of the scFv-GFP in the embryos were driven by the *nos* promoter which was PCR amplified from the *w^1118^*genomic DNA. The *scFv-GFP* sequence was PCR amplified from pHRdSV40-scFv-GCN4-sfGFP-VP64-GB1-NLS (Addgene plasmid # 60904), a gift from Dr. Ron Vale. The PCR fragments were assembled by Gibson assembly, and injected into *y[1] M{vas-int.Dm}ZH-2A w[*]; P{CaryP}attP40* (*attp2, 25C6*) embryos by Core Facility of *Drosophila* Resource and Technology in Core Technology Facility of Center for Excellence in Molecular Cell Science, Chinese Academy of Science. The cloning sequences are listed in “Cloning sequences” section of the supplement text.

### Drosophila stocks

The *w^1118^* (BDSC3605) and *hb* deficient mutant *w^1118^; Df(3R)BSC197/TM6B, Tb[+]* (BDSC9623) came from Bloomington *Drosophila* Stock Center. *w[*]; scFv-sfGFP; Dr/TM3, sb* fly line is constructed in this study. The virgin females of the *w^1118^; P{w[+mC]=GAL4::VP16-nanos.UTR}CG6325[MVD1]* (BDSC4937) were crossed with *w[*]; P{w[+mC]=UAS-GFP-RpL10Ab}BF2* (BDSC42681) from Bloomington *Drosophila* Stock Center for the imaging of the ribosomal protein distribution in the embryos. The virgin females of the *w^1118^; Df(3R)BSC197/TM6B, Tb[+]* were crossed to *hb-SunTag* (*sp/Cyo; hb-SunTag/TM3, sb*) male. The offspring embryos with only one transcription site in each nucleus were selected for quantitative imaging as *hb-def*. *w[*]; sp/Cyo; kni-SunTag/ TM3, sb* fly line was constructed in this study.

### Determination of developmental stages for embryos

The embryos in nc 13 or nc14 were distinguished by the nucleus density. The specific time point of the embryo in nc 14 is determined by the morphology of the nuclei at the mid-sagittal plane based on the reference figures from previous work^31^.

### smFISH and immunofluorescence IF staining

Embryos were collected for 1 h and incubated to the specific developmental stage at 25 °C, and subsequently dechorionated by 4% sodium hypochlorite for 3 min. Following fixation in a 1:1 solution of 4% paraformaldehyde and heptane, the embryos were de-vitellinized in methanol. The methanol is removed through 3 successive washes in PBST (PBS with 0.1% Tween-20 (SIGMA, P1379-500ML)) for 5 min each, followed by an extended 20 min wash. The embryos were then washed three times for 5 min and once for 20 min in hybridization washing buffer (35% Formamide (Invitrogen, 15515026), 20% 20× SSC (Invitrogen, AM9770), and 0.1% Tween-20 in DEPC-treat water (Invitrogen, AM9920)). Subsequently, the hybridization buffer was replaced by smFISH hybridization buffer (35% Formamide, 20% 20× SSC, 1% Dextran sulfate (SIGMA, E008906-10G), 0.1 mg/mL Salmon sperm DNA (Invitrogen, 15632011), 2 mM Ribonucleoside vanadyl complex (NEB, S1402S), 0.2 mg/mL UltraPure BSA (Invitrogen, AM2616), 0.1% Tween-20 in DEPC-treat water) containing oligonucleotide probes specific to target sequences. The embryos were then incubated overnight at 37 °C in the dark. The smFISH probes were designed using the Stellaris Probe Designer https://www.biosearchtech.com/stellaris-designer and synthesized by Stellaris with 3′ fluorophore modifications (probes targeting SunTag sequence: Quasar 570, probes targeting *hb* coding sequence (CDS) region: Qasar 570). The probe sequences for targeting *SunTag* or *hb* CDS (for *hb* mRNA and Hb protein co-staining) are listed in Table S4 and Table S5.

After hybridization, the embryos were washed in pre-warmed (37 °C) hybridization washing buffer in a 3 min-5 min-3min washing sequence, followed by 4 rinses in PBST. The embryos were then stained with DAPI and mounted in Aqua-Poly/Mount (Polysciences, 18606-20), or subjected to IMF staining.

For IF staining, the embryos were washed 3 times for 10 min in PBSX (PBS with 0.1% Triton X-100 (amresco, M143-1L)), followed by 1 h rotation in blocking buffer (western blocking reagent (Roche, 11921673001), 2 mM Ribonucleoside vanadyl complex, 0.1% Triton X-100 in PBS). The embryos were then rotated at R.T. in a primary antibody (GCN4 peptide: mouse anti-GCN4, Novus, NBP2-81273B; or Hb: Abcam, ab197787) solution at a desired dilution in blocking buffer for 3 h. After 3 times for 10 min washing in PBSX and 1 h in blocking buffer, the embryos were further stained by a secondary antibody (Goat anti-Mouse, Alexa Fluor™ Plus 647, Thermofisher, A32728) at a desired dilution in blocking buffer, 2.5 h rotation at R.T. The embryos were then stained with DAPI and mounted.

### smiFISH for the measurement of the end-to-end distance of *hb* mRNA

The 5′ and 3′ ends of *hb* mRNA were targeted by two smiFISH probe sets with Alexa Fluor 594 (AF594) and ATTO 647N labeling, respectively. A typical smiFISH probe consists of a primary probe for RNA targeting and a secondary probe with dye (at both 5′ and 3′ ends of the secondary probe) to recognize the primary probe^47–49^. The probe design and staining processes was conducted as previously described^48^, except the difference in the molecule weight of dextran sulfate (replaced SIGMA, molecular weight 6500–10,000 by SIGMA, E008906-10G). The embryos were collected and fixed as above mentioned and subjected to smiFISH. The corresponding sequences of the primary and secondary probes are listed in Table S6 and Table S7. After smiFISH, the embryos were stained with PicoGreen (yuanye, S33024-100ul) and mounted in SlowFade™ Diamond Antifade mounting medium (Invitrogen, S36967) for STED imaging. For smiFISH and smFISH co-staining for the selection of *hb-def* embryos, the embryos were processed as mentioned in smFISH staining procedure. The annealed smiFISH probes targeting *hb* CDS (Table S8, Alexa Fluor 488 modifications for secondary probe) were directly added to the smFISH hybridization buffer accompanied with smFISH probes. The smiFISH probes for the co-localization control experiment are listed in Table S9.

### mRNA concentration scaling

For the heterozygous *hb-SunTag* fly line, smFISH probes targeting the SunTag sequence detect only the *SunTag*-bearing fraction of the total *hb* mRNA pool transcripted from *hb-SunTag* allele. Multiplying a scale factor of 2 to the detected *N* values (Fig. 1G), the resultant mRNA concentration (*N*_scaled-s_) in the anterior region reaches approximately 600 (Fig. S2), coincides with level detected at similar AP positions in previous work^32^. However, we cannot totally exclude the possible interferences from SunTag insertion or genetic background differences between works

### Imaging setting

The high-resolution imaging of the mRNA (detected by smFISH probes) and corresponding translational signals (GCN4 peptide IMF) were conducted on the Leica TCS SP8 gSTED 3 × (confocal mode), featuring with 63 × oil objective lens (HC PL APO CS2 63 × /1.40 OIL, 1.4 NA), 1.3 zoom factor and 1 AU pinhole size (reference wavelength: 580 nm). An image stack with a vertical range of approximately 15 μm was acquired at the *hb* mRNA profile in each embryo. The Z step size was set to 250 nm, while the XY pixel size was 69 nm. Imaging parameters included 4 times line averaging and a scan speed of 200 Hz. DAPI-stained nuclei, mRNAs (with probes) and translational signals were excited using laser wavelengths of 405 nm (∼3% laser power), 546 nm (∼19% laser power) and 633 nm (∼2.5% laser power), respectively (WLL laser power: 85%). The emission signals were detected in 415–468 (PMT gain: 650), 556–623 (HyD gain: 120, Time Gating: 0.3–11.9 ns, 633 nm reference wavelength) and 643–736 (HyD gain: 85, Time Gating: 0.3–11.5 ns, 633 nm reference wavelength) nm ranges respectively in separate sequences.

For the imaging of smFISH and immunofluorescence co-staining in *w^1118^*embryos, the Leica TCS SP8 gSTED 3 × (confocal mode) was utilized, featuring a 20 × air objective lens (HC PL APO CS2 20 × /0.75 DRY, 0.75 NA) and 0.77 AU pinhole size (reference wavelength: 580 nm). Around mid-sagittal plane, 3 successive images (1024 × 1024 pixels) were acquired in individual embryos with the step size of 0.685 μm. The MIP of each successive image stack was subjected to further quantitative analysis. DAPI-stained nuclei, mRNAs with probes and the antibodies were excited using laser wavelengths of 405 nm (∼2.5% laser power), 546 nm (∼17% laser power) and 633 nm (∼20% laser power), respectively (WLL laser power: 85%), with 3 times line averaging and a scan speed of 200 Hz. The emission signals were detected in 415–468 nm (PMT gain: 600), 558–631 nm (HyD SMD gain: 180, Time Gating: 0.3–11.9 ns, 546 nm reference wavelength) and 643–736 nm (HyD gain: 200, Time Gating: 0.3–12 ns, 546 nm reference wavelength) ranges respectively in separate sequences.

For the imaging of RpL10A-GFP, the embryos were collected and fixed as mentioned above following a DAPI staining. The imaging setting is similar to the *w^1118^* embryos with some adjustments listed below. The pinhole was set as 1 AU (reference wavelength: 580 nm). DAPI and GFP were excited using laser wavelengths of 405 nm (∼6% laser power) and 488 nm (∼3% laser power), respectively (WLL laser power: 85%). The emission signals were detected in 415–467 nm (PMT gain: 800), 498–566 nm (HyD gain: 150) ranges respectively in separate sequences.

In measurement of mRNA end-to-end distance in *w^1118^* embryos by STED, the Leica TCS SP8 gSTED 3 × (STED mode) was utilized, featuring with 100 × oil objective lens (HC PL APO CS2 100 × /1.40 OIL, 1.4 NA), 6.5 zoom factor and 1 AU pinhole size (reference wavelength: 580 nm). The XY pixel size is 17 nm. Imaging parameters included 3 times line averaging and a scan speed of 200 Hz. PicoGreen nucleus staining was excited with 488 nm laser wavelength (∼2%, WLL 85%). The mRNA 5′ and 3′ end labeling was excited by 594nm (∼20% laser power, WLL 85%) and 633 nm (∼20% laser power, WLL 85%), respectively. The depletion laser powers were set as 60% and 70% (775 nm laser line, 0.7760 W) for 5′ and 3′ end labeling respectively. The emission signals were detected in 501–557 nm (PMT gain: 704), 649–732 nm (HyD gain: 80, Time Gating: 0.3–11.0 ns, 633 nm reference wavelength) and 604–637 nm (HyD gain: 80, Time Gating: 0.3–11.8 ns, 633 nm reference wavelength) ranges respectively in separate sequences.

In measurement of mRNA end-to-end distance with detection of anti-GCN4 signals in *hb-SunTag* embryos by STED, the Leica TCS SP8 gSTED 3 × (STED mode) was utilized, featuring with 100 × oil objective lens (HC PL APO CS2 100 × /1.40 OIL, 1.4 NA), and 6.5 zoom factor. 1 AU pinhole size (reference wavelength: 580 nm) for 633 nm channel, 0.8 AU for 594 nm channel, and 0.6 AU pinhole size for 488 nm channel were set. The XY pixel size is 17 nm. Imaging parameters included 3 times line averaging and a scan speed of 400 Hz. Anti-GCN4 staining was excited with 488 nm laser wavelength (∼30%, WLL 85%), depleted by 592 nm depletion laser (∼60% of 592 nm line with 1.0570 W). The mRNA 5′ and 3′ end labeling was excited by 594 nm (∼20% laser power, WLL 85%) and 633 nm (∼20% laser power, WLL 85%), respectively. The depletion laser powers were set as 60% and 70% (775 nm laser line, 0.7750 W) for 5′ and 3′ end labeling respectively. The emission signals were detected in 505–550 nm (HyD gain: 110, Time Gating: 0.3–11.0 ns), 649– 732 nm (HyD gain: 80, Time Gating: 0.3–11.0 ns, 633 nm reference wavelength) and 604–637 nm (HyD gain: 80, Time Gating: 0.3–11.8 ns, 633 nm reference wavelength) ranges respectively in separate sequences.

The localization precision was assessed by fitting the point-spread function to single-molecule objects in each channel. The calculated localization precision (AF594 channel: X direction: 27 ± 4 nm, Y direction: 37 ± 10 nm; ATTO647N channel: X direction: 29 ± 3 nm, Y direction: 28 ± 9 nm) approaches the default spatial resolution of STED microscope. Moreover, the distribution of directions of AF594-ATTO647N pair vectors is approximately uniform in all angles (Fig. S10).

### Imaging processing

Single-molecule images of *hb-SunTag* embryos were processed in a frame-by-frame manner using a custom MATLAB code to detect mRNA and anti-GCN4 spots. Each individual frame of the image stack underwent filtering with a Difference of Gaussian (DoG) filter, utilizing a kernel size of 21 pixels and standard deviation of 1.2 and 2.2 pixels. The image was processed using a Gaussian filter with a standard deviation of 1.5 pixels and a kernel size of 7 pixels, followed by a gray level transformation to facilitate threshold determination. Notably, all transformations mentioned above were exclusively for threshold determination, rather than for the intensity quantification of single molecule objects. An ultimate threshold for the mask segmentation of single molecule objects in each frame was auto-adaptively determined by the intensity value corresponding to the elbow point of mask numbers changing in a series of threshold value applied. To minimize false positives, a three-successive-layer criterion was applied to the mask stack, as described in previous work.^32^ Three-dimension centroids of the single molecule objects within the image stack were further determined by searching for local maxima (with H-maxima transform in MATLAB), adapted from the algorithm established in^50^. As described in previous work^50^, the focal plane image of each three-dimension single molecule object was identified and fitted to a two-dimension gaussian function (eq4) to calculate the integral intensity of this specific object following additional constrains of the parameter values for further excluding noise.^50^ Anti-GCN4 objects were subjected to further quantitative analysis maintaining their intensity values due to the large variation of the object size, while mRNA intensities were converted into absolute counts before further analysis. Besides, the Hb protein is transported to nucleus after translation, in contrast to *hb* mRNA distributed in cytoplasm naturally. It is thus less convincing to assume sufficient single-molecule Hb (anti-GCN4 signals) distribution in cytoplasm for unit intensity determination during count converting. However, the fluorescence intensity is still relatively comparable across positions in individual embryos. The count distribution of integral intensities for all mRNA objects in the image stacks was fitted to a three-term Gaussian function (eq5)^50^. The mean value of the first term was considered as the unit intensity of a single molecule mRNA in individual image stack.

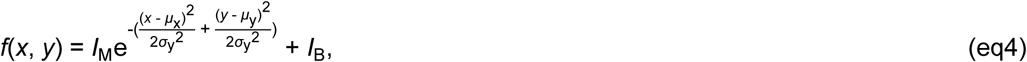

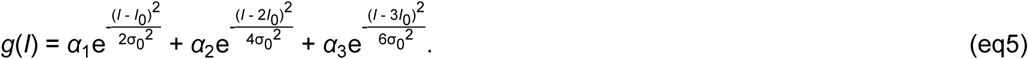

*f*(*x*, *y*) is the intensity at position (*x*, *y*). *I*_M_ and *I*_B_ represent the maximum and baseline intensity of individual single molecule objects. *μ*_x_, *μ*_y_, *σ*_x_ and *σ*_y_ represent the means and standard deviations of Gaussian function at X and Y direction respectively. *g*(*I*) is the count distribution of the integral intensities *I* of mRNA objects. *α*_1_–α_3_ represent 3 amplitude values of 3 Gaussian terms respectively. *I*_0_and *σ*_0_ is the mean and standard deviation of the first term.

After mask segmentation and centroid determination, mRNA and anti-GCN4 signals are regarded as co-localization with Euclidean distance of their centroids smaller than 300 nm.

For the imaging processing of STED images, the end-to-end distance of *hb* mRNA was measured using a custom MATLAB code. The single molecule mRNA objects in each channel were processed separately. Single molecule points were segmented after applying a DoG with kernel size of 21 pixels and standard deviation of 1.2 and 2.2 pixels, following a Gaussian filter with a standard deviation of 1 pixel and kernel size of 5 pixels. A global segmentation threshold was determined by Otsu’s method (graythresh function in MATLAB) with possible manual inspection after gray level transformation. To further remove noise, an open operation was first applied. Then the distribution of the mean intensity for each object was fitted to a Gaussian function (mean: *μ*_G_; standard deviation: *σ*_G_). Objects with mean intensity smaller than *μ*_G_ + 3*σ*_G_ (or *μ*_G_ + 2*σ*_G_, depending on the imaging processing performance), or larger than an upper limit (eg. 10000) were regarded as noise and removed. Besides, a typical mRNA objects should be larger than a threshold in size (AF594 channel: 30 pixels; ATTO 647N channel: 20 pixels). Two points from separate channels were regarded as two ends from one mRNA based on the Euclidean distance of their centroids smaller than 300 nm.

### Statistics

The permutation-based *KW* test (with Bonferroni correction for independent test in individual embryos) and permutation-based Dunn’s test (with Bonferroni correction for pairwise comparisons) were implemented to the data from each embryo respectively due to the relatively large intra embryo variation. The *hb-SunTag* data points from nucleus volume unit in each embryo (background data points in Figure 2 E–G) were binned into 13 groups for *KW* test and the *post-hoc* pairwise comparison. The potential interferences from the shape and variance differences among groups in our data can be properly circumvented by this permutation-based method. The *H* statistic from our observed data were directly compared to the calculated *H* statistic distribution from 10000 permutations of the group labels in this permutation-based *KW* test. The empirical *P*-value is defined as the proportion of the number of calculated permutation *H* statistic with higher value comparing to the *H* statistic from observed data in the permutations. The elevated type I error from the separating *KW* tests of 3 embryos were corrected by Bonferroni correction (adjusted *P*-value). Aligning with the permutation-based statistical procedure, the analytical formula of the denominator of the *z* statistic for *post-hoc* pairwise comparison Dunn’s test was replaced by standard deviation calculated from our data, and the distribution of *z* statistic is determined by 10000 permutations of the group labels. The empirical *P*-value is defined as the proportion of the *z* statistic from permutation with higher absolute value comparing to the absolute value of the observed *z* statistic from our data. The Bonferroni correction also implemented to these pairwise *P*-values (adjusted *P*-values). The same statistical procedures were applied to *TR* from STED-based *hb* mRNA end-to-end distance measurement in each embryo separately. The permutation-based *KW* test was also implemented to the translating mRNA end-to-end distance from the STED imaging.

### Modeling and calculation

The relationships between *TR* and *N* were established by plotting the *TR* changes as *N* increases, based on the parameters from the model fitting of eq1.

The parameters *TR*_offset_, *TR*_b_ and *C*_T_ of the mRNA concentration–dependent model (eq1) were optimized by lsqnonlin in MATLAB with 10000 initial parameter values from Latin hypercubic sampling. The sampling ranges were 0–10, 0–10 and 0–200 respectively. The model fitting results are listed in Table S2.

The constant model and mRNA concentration–dependent model were fitted to Hb protein profile in 0.25–0.55 EL at ∼12 min of nc 14, with the input mRNA and initial Hb protein profile at ∼6 min of nc 14 (Data source: reference 6). The parameters *k*_offset_, *k*_b_ and *C*_T_ of the mRNA concentration–dependent translational model and *k* of constant model were optimized by lsqnonlin in MATLAB with 10000 initial parameter values from Latin hypercubic sampling, respectively. The sampling ranges were 0–10, 0–10, 0–200 and 0–1 for*k*_offset_, *k*_b_, *C*_T_ and *k*. Parameter *D* = 0.005 (EL^2^ / min) represents the diffusion constant of the protein Hb adapted from previous works ^5,51,52^. The Hb protein degradation rate (min^−1^) was a given value (0.005, 0.010, 0.015, 0.020, 0.025) which was consistent in two models. The model fitting results are listed in Table S3.

We simulated the protein profile from two models to later time points (∼24 to 48 min of nc14) with parameter values from their model fitting at ∼12 min of nc 14 with corresponding protein degradation rates. Protein values in the 0.27–0.33 EL were averaged for both experimental data and model prediction values and used for RSS calculations.

## DATA AVAILABILITY

Further information and requests for resources and reagents should be directed to and will be fulfilled by the lead contact, Feng Liu.

All materials other than fly line and probes are commercially available. Fly lines and probes can be provided upon request in accordance with Chinese regulations.

Data related to figures have been deposited at Mendeley Data and are publicly available as of the date of publication at https://data.mendeley.com/preview/rn3cy4rxtv?a=d146353b-9133-42e7-9d57-c274c49c39cf, or cited to previous publication.^6^ All produced data in this paper will be shared by the lead contact upon request.

Image processing codes and statistical analysis codes have been deposited at Mendeley Data and are publicly available as of the date of publication at https://data.mendeley.com/preview/rn3cy4rxtv?a=d146353b-9133-42e7-9d57-c274c49c39cf.

## ACKNOWLEDGMENTS

We thank Martin Gruebele for careful reading and discussion. We thank Yihan Lin for providing the facilities required for vector construction. This project is supported by the National Natural Science Foundation of China 32271293. The numerical calculation was performed on the High-Performance Computing Platform of the Center for Life Sciences, Peking University.

## ETHICS DECLARATION

The authors declare no competing interests.

