## Supplement text, figures, and tables. for "mRNA concentration–dependent translation enables rapid and sharp patterning in resource-constrained *Drosophila* embryos"

### Cloning sequences

#### *hb-SunTag* sequence:

ccactgcatggtccccatgcccccacccctgccccctggcccatgcccctgcccggcgtaatattaattttacacttggcctttagttt
ggctttgttgctgttgttgggcatggcacaaaaaagcccagacgaaaggcgaaaattctctttgtttcatagtgcgcacacacacta
acacacttgccgagcaattaattttggctttctttaaaaatatccacaaaagcgaatttaaataaatatgctaagcttcattttgtgtggt
gcactttctgtttcctgaaccatcaattatgcctaagtattgtagatattttagctgccagatagcacagcaccatcctcccataataat
attccgtaaatgccccttttcccgttttgcgtttttaataatatttacttgaaagcacaaacaattagccaaaaatgcagcaactgcac
aatttttcagtgtgaaattggaaatggaaaaaatataggcaacaagcaattttaatgcgagaattattagaaaaactacgcaaatc
aaagtgaaatgtctggcggaaaattgttgtcaggaaaatgtttttcaaatgggtgtgtaataattatactgcacatattatgcatatagt
ttagttggtccttagagttttcccgcaggtgtaagcagttctgatccgttaatttagttaagtcccgcaatcctttttactttttattattaacta
cgaaactgcccacgctagctgcctactcctgctgtcgactcctgaccaacgtaatccccatagaaaaccggtggaaaattcgca
gctcgctgctaagctggccatccgctaagctcccggatcatccaaatccaagtgcgcataatttttttgtttctgctctaatccagaatg
gatcaagagcgcaatcctcaatccgcgatccgtgatcctcgattcccgaccgatccgcgacctgtacctgacttcccgtcacctct
gcccatctaatcccttgacgcgtgcatccgtctacctgagcgatatataaactaatgcctgttgcaattgttcagtcagtcacgagttt
gttaccactgcgacaacacaacagaagcagcaccaataatatacttgcaaatccttacgaaaatcccgacaaatttggaatata
cttcgatacaatcgcaatcatacgcactgagcggccacgaaacggtaggatattgttagccattaccaagtgtctccattttgaaca
caaaatcactcaaatcgccttcagggggtgggtgccgcccagccacccctgacgtatttttttgttaggggtggtgccgcaagcaca
ccaaaaaaagagaaaaaaaaaaataaaagcgaggaaaaataaaatgaaaaacaagcggaaaaaaagaggaaaaaact
cgacgcaggcgcagtgcatgaatgaataaatgaatatgcccactaaccccactctctctgttttcttatccattacagccgtctagag
ccgccaagatggaagaacttttgagcaagaattatcatcttgagaacgaagtggctcgtcttaagaaaggttctggcagtggaga
agaactgctttcaaagaattaccacctggaaaatgaggtagctagactgaaaaaggggagcggaagtggggaggagttgctg
agcaaaaattatcatttggagaacgaagtagcacgactaaagaaagggtccggatcgggtgaggagttactctcgaaaaattat
catctcgaaaacgaagtggctcggctaaaaaagggcagtggttctggagaagagctattatctaaaaactaccacctcgaaaat
gaggtggcacgcttaaaaaagggaagtggcagtggtgaagagctactatccaagaattatcatcttgagaacgaggtagcgcg
tttgaagaagggttccggctcaggagaggaactgctctcgaagaactatcatcttgaaaatgaggtcgctcgattaaaaaaggg
atcgggcagtggtgaggaactactttcaaagaattaccacctcgaaaacgaagtagctcgattaaagaaaggttcagggtcgg
gtgaagaattactgagtaaaaattatcatctggaaaatgaggtagcgagactaaaaaaggggagtggttctggcgaagagttgc
tatcgaaaaattatcatcttgagaacgaagttgctaggctcaaaaagggctcaggctcaggcgaggagttgctctcgaaaaacta
ccacttggaaaatgaggtcgcgaggttgaaaaaggggagcgggtcgggcgaggagttattgagcaaaaactatcatttagaga
acgaagtcgcgcgcttaaagaaaggctcgggctcgggcgaagaactcttatcgaagaactaccacctcgaaaatgaggtcgc
caggttgaaaaagggcagtggcagcggggaggaactcttgagcaagaactaccacttggagaatgaggtcgcgagattgaa
gaaagggtcggggagcggcgaggaattgctcagcaagaattatcatttggagaacgaagtcgccaggctcaagaaaggctc
ggggtcgggggaggaattgttgagtaaaaactaccacttggaaaatgaagtcgccaggctcaaaaaagggagtgggagcgg
cgaagagttattgagcaaaaattaccacttggagaacgaagtggcaaggctcaagaaagggagcggcagcggggaggagc
tcttatcgaagaactaccacttagagaatgaagtcgcccgcttgaagaaaggctcggggagcggggaagagctcttgagcaag
aactaccacttggaaaatgaggtggcgcgcttgaagaaagggagcgggagcggggaagagttactatctaagaattatcatct
cgagaacgaggtggctcgactaaagaagggctccggcagtggggaggaactcctgtcgaagaactatcatcttgaaaatgag
gttgcaagacttaaaaaggggtccggatcaggtgaggaactactcagtaagaattaccacctggaaaacgaagttgcacgtttg
aagaaaggatcaggatcaggcgaagaactgctctcaaaagattatcatttggaaaatgaggttgcacgtttaaaaaagggaag
tggcagtggtgaggaacttctgtcgaaaaattatcatctcgagaatgaagtagcccgacttaaaaagggtggggaggaggag
gaagatctcagaactgggagacgacagccacgaccaactacgagcagcacaacgcctggtacaacagcatgttcgcggca
aatatcaaacaggagccaggtcatcatctcgacgggaatagcgtggcaagcagtccgcgccaatcgcccattccctcgaccaa
tcacctggaacagttcctcaagcagcagcagcagcagcttcagcagcaacccatggataccctgtgcgccatgaccccatcac
ccagccaaaacgatcaaaacagcctgcagcattacgatgctaacttgcagcaacagttgctgcagcaacagcagtaccagca
gcatttccaggcagcccagcagcaacatcatcaccatcaccatctgatgggtggattcaatccgctgacgccacctggtctgccc
aatcccatgcagcacttctatggcggcaatctgcgacccagtccgcagcccacgcccacatctgcctccacaattgcgcccgttg
cagttgccactggcagcagcgagaagttgcaggcactaacaccacccatggatgtcacaccgcctaagtcgccggccaagtc
gagtcagtcgaatattgagccggagaaggagcacgatcagatgtcgaactccagcgaggacatgaagtacatggccgagtcc
gaggacgatgataccaacatccggatgcccatctacaattcgcacggcaagatgaagaactacaagtgcaagacctgcggcg
tggtggccatcaccaaggtggacttctgggcgcacacccgcacccacatgaaaccagacaagatcctgcagtgcccgaagtg
cccgttcgtcaccgagttcaagcaccacttggagtaccatatccggaagcacaagaaccaaaagcccttccagtgcgacaaat

gcagctacacgtgtgtcaacaaatccatgctaaactcgcaccgcaagtcgcacagttctgtgtatcagtaccgttgtgcggattgtg
attacgccaccaagtattgccacagcttcaagctgcatctgcgcaagtatggtcacaagcccggcatggttttggacgaggatgg
caccccgaatccctcgttggtcatcgatgtttacggcacgcgtcgtggtccgaagagcaagaatggtggaccgattgccagtgga
ggaagtggcagcggcagccggaagtcaaatgttgcagctgtcgctccgcagcaacagcaatctcagccagctcagccagtcg
ccacatctcagctgagtgccgccctgcaaggattccctctggttcaaggcaactccgctcctccggcggcatctccagtgctcccg
ctgcccgcctctcctgccaagagtgtggccagtgtggaacagacgcccagcttgcccagtccagccaatcttctgcctcctctggc
cagccttctgcagcagaaccgcaacatggccttcttcccctactggaacctcaatctccagatgctggccgcccaa

***kni-SunTag* sequence:**
atgagcagctgttaaaaatgaagtcaatagcttagtcaatgtggtgtgtgtgtgtgtgtggtgttcgagtgagtgagaaatccagccgcc
cttagcacgcgagtatctttaataaataaacgaataacgaataatatcagggccatgcaaatagcctgattacagggaactcaaaatcg
agagagagagagagactgagcgtgagatctgaatgagtgagtgagtgttttctattcattcaacaacagagcgttaacattctgctaaca
tttcgctcgagtggggttcgaactcaatgcgcatgtgtgcgtgctcgatcgctctctcactcgatccgagtcttaaaggtggtggtttcagccg
tgatttatcagagagctggggctgaaaactggtaagtttgctttggtgtgagtgcgagtacataagccaaagagttcggccagtaggcga
aacacagaaacgcttcttgccagtcgaacctctcagccagaaatcagtcgtcagctagtagccgaagtgtcagttacacacatcgagg
attcccaaaccgcgttgtttcggcgatcaaaaaccatacatcacatcaaatcgtatccaaaattataatcaaagtgttgaaaacctgaatc
actcagactgatcgaaaagtgcttgcaaaccgaaaacaaaacaaaaaaaaaaaaaaataacagacacaaactgccaaagtgaga
aaacgtgcagcaacgatcgcgaatcgctgcaattaaaaagtcgtcataaacctagtgcccatccaacaaaaaaaagcaaaaaggta
agttctgagcgagggagacaacatgagatcaaagtcaattcacggttcttgtttttgggtcgcggccgtcggcaaattttacgatcccttgtg
cactacttttattttttactgtttcacgcgaaaaactaacggccacatttccatcttttccttatttttttgcgtcccgaggcagcgggaaaaaaat
cacaattttcataagccgaattttctatttttgtttgactcgggaaaaatcggcgtgagtttttatggccgaacgtcaaggtctggactgctgct
gcattttttagggaacactattttcgcagcactcgtaatgacttcgaataaaaaagaaaatctacctgagttttatgactggacagcgcga
aaaaaatgaaaatgaacgcatagggttgcataatccggcagcttaagtttttggcctgttgtgatcataaaaaacgcccatcctgtctagt
ttttcccagtttcctatatatatcctggcccgcctagagctcagcatcagttgctcagcagcattccaagcgaacagatcatacagcagatc
ctcacagcgatcgtgagaaaagcattcaaaattccaacaaatatcattccaaaatggtgttcaacttggttagtgtccagagccgttcgcc
tacttgtggattacacatatacctctcgaccagtggattaaacccttactaacgcgTgatttctttacaatcttccagatggaagaacttttga
gcaagaattatcatcttgagaacgaagtggctcgtcttaagaaaggttctggcagtggagaagaactgctttcaaagaattaccacctgg
aaaatgaggtagctagactgaaaaaggggagcggaagtggggaggagttgctgagcaaaaattatcatttggagaacgaagtagc
acgactaaagaaagggtccggatcgggtgaggagttactctcgaaaaattatcatctcgaaaacgaagtggctcggctaaaaaaggg
cagtggttctggagaagagctattatctaaaaactaccacctcgaaaatgaggtggcacgcttaaaaaagggaagtggcagtggtgaa
gagctactatccaagaattatcatcttgagaacgaggtagcgcgtttgaagaagggttccggctcaggagaggaactgctctcgaaga
actatcatcttgaaaatgaggtcgctcgattaaaaaagggatcgggcagtggtgaggaactactttcaaagaattaccacctcgaaaac
gaagtagctcgattaaagaaaggttcagggtcgggtgaagaattactgagtaaaaattatcatctggaaaatgaggtagcgagactaa
aaaaggggagtggttctggcgaagagttgctatcgaaaaattatcatcttgagaacgaagttgctaggctcaaaaagggctcaggctca
ggcgaggagttgctctcgaaaaactaccacttggaaaatgaggtcgcgaggttgaaaaaggggagcgggtcgggcgaggagttatt
gagcaaaaactatcatttagagaacgaagtcgcgcgcttaaagaaaggctcgggctcgggcgaagaactcttatcgaagaactacca
cctcgaaaatgaggtcgccaggttgaaaaagggcagtggcagcggggaggaactcttgagcaagaactaccacttggagaatgag
gtcgcgagattgaagaaagggtcggggagcggcgaggaattgctcagcaagaattatcatttggagaacgaagtcgccaggctcaa
gaaaggctcggggtcgggggaggaattgttgagtaaaaactaccacttggaaaatgaagtcgccaggctcaaaaaagggagtggg
agcggcgaagagttattgagcaaaaattaccacttggagaacgaagtggcaaggctcaagaaagggagcggcagcggggagga
gctcttatcgaagaactaccacttagagaatgaagtcgcccgcttgaagaaaggctcggggagcggggaagagctcttgagcaagaa
ctaccacttggaaaatgaggtggcgcgcttgaagaaagggagcgggagcggggaagagttactatctaagaattatcatctcgagaa
cgaggtggctcgactaaagaagggctccggcagtggggaggaactcctgtcgaagaactatcatcttgaaaatgaggttgcaagactt
aaaaaggggtccggatcaggtgaggaactactcagtaagaattaccacctggaaaacgaagttgcacgtttgaagaaaggatcagg
atcaggcgaagaactgctctcaaaagattatcatttggaaaatgaggttgcacgtttaaaaaagggaagtggcagtggtgaggaacttc
tgtcgaaaaattatcatctcgagaatgaagtagcccgacttaaaaagggtgggggaggaggaggaagatctaaccagacatgcaaa
gtgtgcggtgagccggcggcgggcttccattttggcgccttcacctgcgagggctgcaaggtaagttgtgtctcaagaaatccattgaca
aataaattagcagaacgtaacccccccaaggggttagttttagaaatgttcgaggaacaggccatcggcgattcaaatcgttgtaccattg
gccttaagttcttgaatgaatttctgctcttcttttgctaatgagttgcataccacataactaagccacattcgtccttcccttcgccattgcagtcc
ttctttggccgctcttacaacaacatcagcaccatcagcgagtgcaagaacgagggcaagtgcatcatcgacaagaagaaccgcacc
acctgcaaggcgtgccgcttgaggaagtgctacaacgtgggcatgtcgaagggggggatcccgctacggacgtcgctccaactggttca
agatccattgtctgctgcaggagcacgaacaggccgccgcagcggcgggcaaggcgcctccattagcgggtggcgtatcggtgggtg
gtgccccgtcggcctcttccccggtgggctcgccacacactcccggatttggggacatggccgcccatttgcaccaccatcatcagcag
cagcagcagcagcaggtgccgcgtcatccacatatgcctctgctgggctatcccagctatctgtccgacccatccgccgccctgcccttct
tcagcatgatgggcggtgtaccgcaccagtcgcccttccagctgcccccacacctcctcttcccaggctaccatgcaagtgctgccgctg
cagcggcttctgctgccgatgccgcttaccggcaggagatgtacaagcaccgccagagcgtggattccgttgagtcgcagaaccgcttt

agtcccgccagccagccaccagtggtgcagcccacctcctcggcccgccagtcgcccatcgatgtctgcctggaggaggatgttcact
ccgtgcacagccatcagtcgtccgcaagcctcctgcatcccattgccatccgagccacgccaaccactccgactagcagcagcccgct
gagttttgcggccaagatgcagagcttgtcgcccgtttcggtttgctccattggcggcgaaaccaccagcgttgtaccagtgcatcctccca
ccgtttccgctcaagaaggacccatggatctgagcatgaagacctcgcggagctccgtgcacagcttcaacgacagcggctccgagg
atcaagaagtggaggtggctccgcgccggaagttctaccaactggaggccgagtgcctgaccaccagcagcagcagttcctcccact
ccgccgcccactcaccgaacaccaccaccgcccatgcggaagtcaagcggcagaagctaggtggtgcagaggccacccacttcgg
tggcttcgcggtggcccacaatgcggctagtgccatgaggggaatattcgtgtgtgtctaagtacacggcgaaaaaaccaagtgggag
gagtc

***scFv-sfGFP sequence:***
aaggagaaaaaatcattgaaagcttcgaccgttttaacctcgaaatatgcacatgtaaggacggatgtgagcgaacgccagtg
atgaccgggatcagaggtaacctaccatggtggggattaggtgaccgttcgcaggtagtttgatcggagcgaatgttcggggggt
ctggcgtcagaggctctaaactttatgtaattcctgccgcgaaacacgcacgtatcaagcagtcagctgttctcttcgttcagcgcgc
gccggtgttgcaaaacgagcgctcttcgccggcggtggctcgtgcgatagttcgttttgtcggtaatccgatgttgccgcgccgatat
catgtgatgttgtcacagtgcgcgaaattcgaatggtggtgtgcagtgattgtgttgtgacggcgagtggcgcgtgtgggtgcttagtt
ttgggagatgttttcgtattttttgttgataactcaggctttgttgctgtgttgtagtactattttccattgcgcggtgtccagcttttaattagtg
gcacatattcttagcaagtaaaaattattttgcatactattaaatttcttataaattattttctaaaattaagtttaccttttcaattttactaaa
aatatcgatatatttattatcgctggaaaactacattattccacctctaagcaagaaccgttagttggcgcgtagctttaccacaaaat
tcctggaattgccgtacgcttcgcagttgtttcaagttgtctaagggacatacgattttttttgcctctgcgtcacgattttaacccaaaag
cgagtttagttacatgtacattattattagataaagaagtatcgcgaatacttcagttgaataaactgtgcttggtttttgggtgaggattt
gtggaaagtagagtgcgcgataaccgtaactttcgacccggattttcgccgccaccatggccccgacatcgtgatgacccaga
gccccagcagcctgagcgccagcgtgggcgaccgcgtgaccatcacctgccgcagcagcaccggcgccgtgaccaccagc
aactacgccagctgggtgcaggagaagcccggcaagctgttcaagggcctgatcggcggcaccaacaaccgcgccccccgg
cgtgcccagccgcttcagcggcagcctgatcggcgacaaggccaccctgaccatcagcagcctgcagcccgaggacttcgcc
acctacttctgcgccctgtggtacagcaaccactgggtgttcggccagggcaccaaggtggagctgaagcgcggcggcggcgg
cagcggcggcggcggcagcggcggcggcggcagcagcggcggcggcagcgaggtgaagctgctggagagcggcggcg
gcctggtgcagcccggcggcagcctgaagctgagctgcgccgtgagcggcttcagcctgaccgactacggcgtgaactgggtg
cgccaggcccccggccgcggcctggagtggatcggcgtgatctggggcgacggcatcaccgactacaacagcgccctgaag
gaccgcttcatcatcagcaaggacaacggcaagaacaccgtgtacctgcagatgagcaaggtgcgcagcgacgacaccgcc
ctgtactactgcgtgaccggcctgttcgactactggggccagggcaccctggtgaccgtgagcagctacccatacgatgttccag
attacgctggtggaggcggaggttctggggaggaggtagtggcggtggtggttcaggaggcggcggaagcttggatccaggt
ggaggtggaagcggtagcaaaggagaagaacttttcactggagttgtcccaattcttgttgaattagatggtgatgttaatgggcac
aaattttctgtccgtggagagggtgaaggtgatgctacaaacggaaaactcacccttaaatttatttgcactactggaaaactacct
gttccgtggccaacacttgtcactactctgacctatggtgttcaatgcttttcccgttatccggatcacatgaaacggcatgactttttca
agagtgccatgcccgaaggttatgtacaggaacgcactatatctttcaaagatgacgggacctacaagacgcgtgctgaagtca
agtttgaaggtgatacccttgttaatcgtatcgagttaaagggtattgattttaaagaagatggaaacattcttggacacaaactcga
gtacaactttaactcacacaatgtatacatcacggcagacaaacaaaagaatggaatcaaagctaacttcaaaattcgccacaa
cgttgaagatggttccgttcaactagcagaccattatcaacaaaatactccaattggcgatggccctgtccttttaccagacaacca
ttacctgtcgacacaatctgtcctttcgaaagatcccaacgaaaagcgtgaccacatggtccttcttgagtttgtaactgctgctggg
attacacatggcatggatgagctctacaaaggtggaggtcggaccgaagagtacaagcttatcctgaacggtaaaaccctgaa
aggtgaaaccaccaccgaagctgttgacgctgctaccgcggaaaaagttttcaaacagtacgctaacgacaacggtgttgacg
gtgaatggacctacgacgacgctaccaaaaccttcacggtaaccgaaggtggtggtagcggtggtggtactagtcccaagaag
aagcgcaaggtgtaaaccgcggcacactaagcgtcgcgccacttcaacgctcgatgggagcgtcattggtgggcggggtaac
cgtcgaaatcagtgtttacgcttccaatcgcaacaaaaaattcactgcaacactgaaaagcatacgaaaacgatgaagattgta
cgagaaaccataaagtattttatccacaaagacacgtatagcagaaaagccaagttaactcggcgataagttgtgtacacaaga
ataaaatcggccagattcagtgttgtcagaaataagaaaccccactatgtttttctttgcctttttcttctcccagcgatcattcatttcgt
ggtgaaagaacggggtcattgcacggagtttcgactgcgggaaagcagagctgccgttcacttcgtctataattagcgctttctattt
tccccgattcgggccgctgctgcgcttttccgcctgctgtttgtggcaagtgtagcagcaggctgtgcacgcagtgtggcatgcactt
ggctttccaccgttggtatcgattctctgggacgatgagtcattcctttcggggccacagcataatcgttgccagctcaccgaaatgg
tgacttcatttcttaactgccgtcaagcatgcgattgtacatacatacatatttatatatgtacatatttatgtgactatggtaggtcgatat
aatagcaatcaacgcaagcaaatgtgtcagtcctgcttacaggaacgattctatttagtaattttcgttgtataaagtaattatgtatgt
atgtaagccccataaatctgaaacaattaggcaaaaccatgcgaagctg

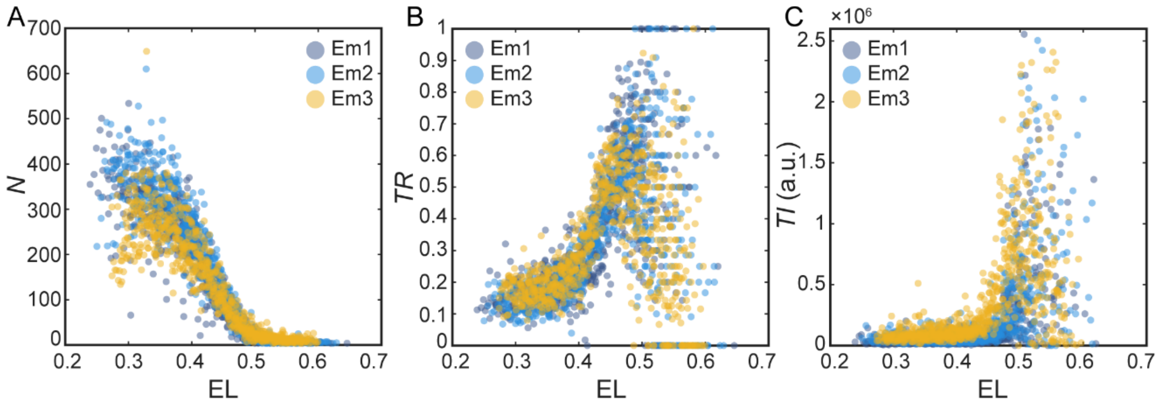


**Fig. S1.** Incorrect spatial heterogeneities of *hb* translation detection in *hb-SunTag* embryos with *in vivo* scFv-GFP detector expression (*w[*]; scFv-sfGFP; hb-SunTag/TM2*). (A, B, C) Spatial distributions of (A) mRNA concentration *N*, (B) *TR* and (C) *TI* along the AP axis from three replicates. The AP axis is normalized to embryo length (EL) with 0 and 1 representing the anterior and posterior pole, respectively. Points represent the values in nucleus volume unit.

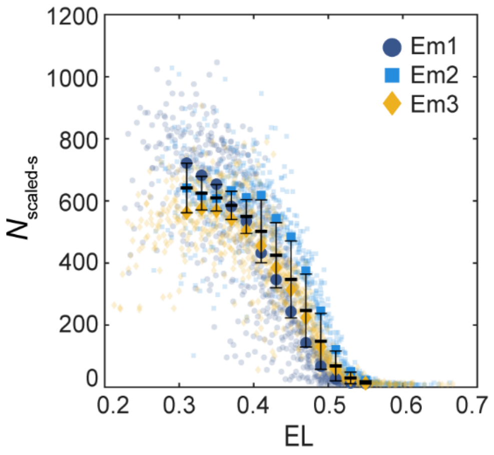


**Fig. S2.** Scaled *hb-SunTag* mRNA concentration along the AP axis. Zygotic transcripts in heterozygous *hb-SunTag* (*w[*]; sp/Cyo; hb-SunTag/ TM6B, Tb[+]*) embryos are imaged.

| EL | 0.31 | 0.33 | 0.35 | 0.37 | 0.39 | 0.41 | 0.43 | 0.45 | 0.47 | 0.49 | 0.51 | 0.53 | 0.55 |
|---|---|---|---|---|---|---|---|---|---|---|---|---|---|
| 0.31 | | | | | | | | | | | | | |
| 0.33 | 1.000 | | | | | | | | | | | | |
| 0.35 | 1.000 | 1.000 | | | | | | | | | | | |
| 0.37 | 1.000 | 1.000 | 1.000 | | | | | | | | | | |
| 0.39 | 1.000 | 1.000 | 1.000 | 1.000 | | | | | | | | | |
| 0.41 | 0.109 | 0.343 | 1.000 | 1.000 | 1.000 | | | | | | | | |
| 0.43 | 0.016 | 0.008 | 0.203 | 0.070 | 0.523 | 1.000 | | | | | | | |
| 0.45 | 0.008 | 0.008 | 0.008 | 0.008 | 0.016 | 0.780 | 1.000 | | | | | | |
| 0.47 | 0.023 | 0.023 | 0.655 | 0.070 | 1.000 | 1.000 | 1.000 | 1.000 | | | | | |
| 0.49 | 0.179 | 0.694 | 1.000 | 1.000 | 1.000 | 1.000 | 1.000 | 0.335 | 1.000 | | | | |
| 0.51 | 1.000 | 1.000 | 1.000 | 1.000 | 1.000 | 0.031 | 0.008 | 0.008 | 0.016 | 0.055 | | | |
| 0.53 | 1.000 | 0.796 | 0.016 | 0.047 | 0.008 | 0.008 | 0.008 | 0.008 | 0.008 | 0.008 | 1.000 | | |
| 0.55 | 0.172 | 0.016 | 0.008 | 0.008 | 0.008 | 0.008 | 0.008 | 0.008 | 0.008 | 0.008 | 0.016 | 1.000 | |

**Fig. S3.** The adjusted *P*-values of the *post-hoc* pairwise comparison Dunn's test for *hb-SunTag* Em1. The *P*-values are corrected by Bonferroni correction. The data points with the *P*-values less than 0.05 are marked as red. The AP axis is normalized to embryo length (EL) with 0 and 1 representing the anterior and posterior pole, respectively.

| EL | 0.31 | 0.33 | 0.35 | 0.37 | 0.39 | 0.41 | 0.43 | 0.45 | 0.47 | 0.49 | 0.51 | 0.53 | 0.55 |
|---|---|---|---|---|---|---|---|---|---|---|---|---|---|
| 0.31 | | | | | | | | | | | | | |
| 0.33 | 1.000 | | | | | | | | | | | | |
| 0.35 | 1.000 | 1.000 | | | | | | | | | | | |
| 0.37 | 1.000 | 1.000 | 1.000 | | | | | | | | | | |
| 0.39 | 1.000 | 1.000 | 1.000 | 1.000 | | | | | | | | | |
| 0.41 | 1.000 | 1.000 | 1.000 | 1.000 | 1.000 | | | | | | | | |
| 0.43 | 1.000 | 1.000 | 1.000 | 1.000 | 1.000 | 1.000 | | | | | | | |
| 0.45 | 1.000 | 1.000 | 1.000 | 1.000 | 1.000 | 1.000 | 1.000 | | | | | | |
| 0.47 | 1.000 | 1.000 | 1.000 | 1.000 | 1.000 | 1.000 | 1.000 | 1.000 | | | | | |
| 0.49 | 1.000 | 1.000 | 1.000 | 1.000 | 1.000 | 1.000 | 1.000 | 1.000 | 1.000 | | | | |
| 0.51 | 1.000 | 1.000 | 1.000 | 1.000 | 1.000 | 1.000 | 1.000 | 1.000 | 1.000 | 1.000 | | | |
| 0.53 | 0.016 | 0.008 | 0.008 | 0.008 | 0.008 | 0.008 | 0.008 | 0.008 | 0.008 | 0.008 | 0.016 | | |
| 0.55 | 0.008 | 0.008 | 0.008 | 0.008 | 0.008 | 0.008 | 0.008 | 0.008 | 0.008 | 0.008 | 0.008 | 1.000 | |

**Fig. S4.** The adjusted *P*-values of the *post-hoc* pairwise comparison Dunn's test for *hb-SunTag* Em2. The *P*-values are corrected by Bonferroni correction. The data points with the *P*-values less than 0.05 are marked as red. The AP axis is normalized to embryo length (EL) with 0 and 1 representing the anterior and posterior pole, respectively.

| EL | 0.31 | 0.33 | 0.35 | 0.37 | 0.39 | 0.41 | 0.43 | 0.45 | 0.47 | 0.49 | 0.51 | 0.53 | 0.55 |
|---|---|---|---|---|---|---|---|---|---|---|---|---|---|
| 0.31 | | | | | | | | | | | | | |
| 0.33 | 1.000 | | | | | | | | | | | | |
| 0.35 | 1.000 | 1.000 | | | | | | | | | | | |
| 0.37 | 1.000 | 1.000 | 1.000 | | | | | | | | | | |
| 0.39 | 1.000 | 1.000 | 1.000 | 1.000 | | | | | | | | | |
| 0.41 | 1.000 | 1.000 | 1.000 | 1.000 | 1.000 | | | | | | | | |
| 0.43 | 1.000 | 1.000 | 1.000 | 1.000 | 1.000 | 1.000 | | | | | | | |
| 0.45 | 1.000 | 1.000 | 1.000 | 1.000 | 1.000 | 1.000 | 1.000 | | | | | | |
| 0.47 | 1.000 | 1.000 | 1.000 | 1.000 | 1.000 | 1.000 | 1.000 | 1.000 | | | | | |
| 0.49 | 1.000 | 1.000 | 1.000 | 1.000 | 1.000 | 1.000 | 1.000 | 1.000 | 1.000 | | | | |
| 0.51 | 1.000 | 1.000 | 1.000 | 1.000 | 1.000 | 1.000 | 1.000 | 1.000 | 1.000 | 1.000 | | | |
| 0.53 | 0.109 | 0.055 | 0.086 | 0.008 | 0.008 | 0.008 | 0.008 | 0.008 | 0.008 | 0.008 | 0.008 | | |
| 0.55 | 0.008 | 0.008 | 0.008 | 0.008 | 0.008 | 0.008 | 0.008 | 0.008 | 0.008 | 0.008 | 0.008 | 0.008 | |

**Fig. S5.** The adjusted *P*-values of the *post-hoc* pairwise comparison Dunn's test for *hb-SunTag* Em3. The *P*-values are corrected by Bonferroni correction. The data points with the *P*-values less than 0.05 are marked as red. The AP axis is normalized to embryo length (EL) with 0 and 1 representing the anterior and posterior pole, respectively.

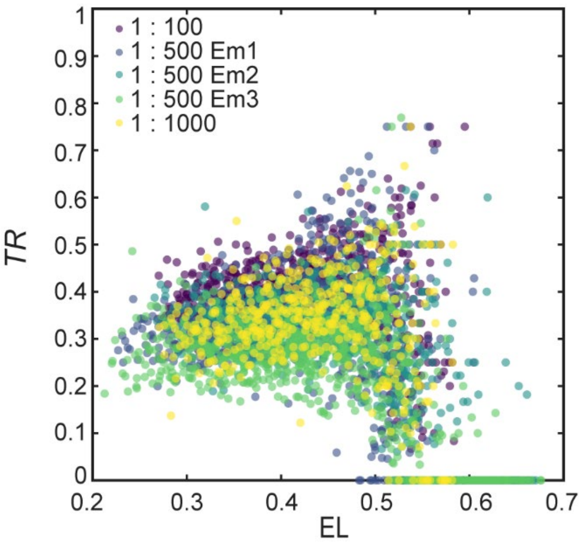


**Fig. S6.** *TR* of *hb* is consistent in *hb-SunTag* embryos stained with dilution ratios of antibodies varying from 1:100 to 1:1000 for SunTag detection. Points represent the values in nucleus volume unit. The AP axis is normalized to embryo length (EL) with 0 and 1 representing the anterior and posterior pole, respectively.

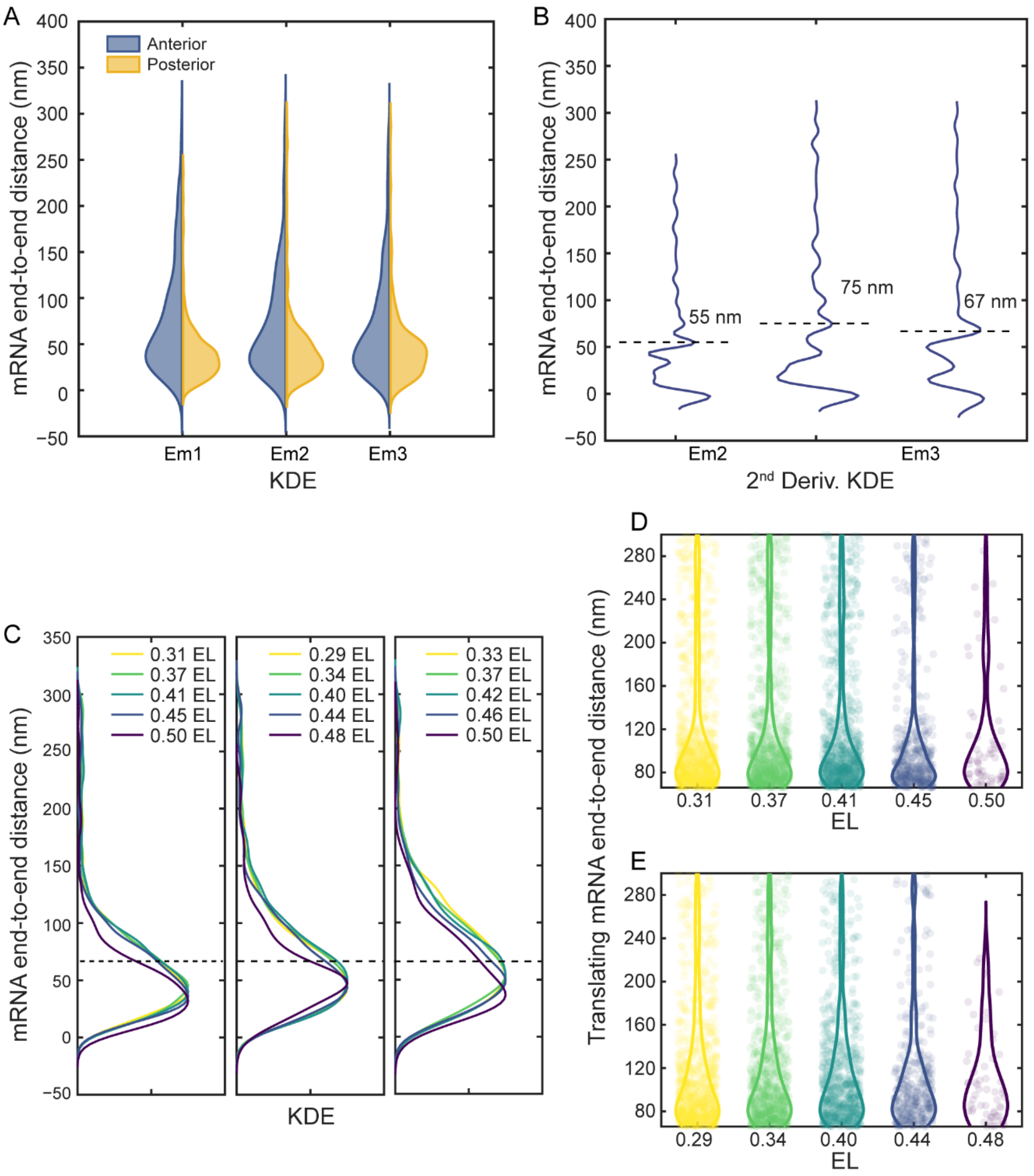


**Fig. S7.** The measurement of the mRNA end-to-end distance reveals spatial heterogeneities of the *hb* mRNA translation during embryogenesis in replicates. (A) The kernel density estimates (KDE) of the *hb* mRNA end-to-end distance in the anterior (sample size n = 513 for Em1, n = 559 for Em2, n = 508 for Em3) and posterior (n = 328 for Em1, n = 432 for Em2, n = 206 for Em3) regions of AP axis for two replicates in nc 6–8. (B) The corresponding second derivative ($2^{nd}$ Deriv.) of KDEs of the *hb* mRNA end-to-end distance in the posterior region of AP axis. Dashed lines annotate the determined distance thresholds for each embryo. (C) KDEs of *hb* mRNA end-to-end distance from 5 positions along AP axis of 3 replicates in nc 14. The AP axis is normalized to embryo length (EL) with 0 and 1 representing the anterior and posterior pole, respectively. Dashed line annotates the determined distance threshold (66 nm). (D, E) Violin plots for translating mRNA end-to-end distances for *hb* mRNAs (background points) from 5 AP positions in the corresponding embryos in (C). (D: corresponding to left panel in C, n = 915, 920, 881, 485, 62. Adjusted *P*-value of the permutation-based *KW* test with Bonferroni correction: 0.0690. E: corresponding to middle panel in C, n = 871, 839, 732, 396, 57. Adjusted *P*-value of the permutation-based *KW* test with Bonferroni correction: 1.000.).

A

| EL | 0.31 | 0.37 | 0.41 | 0.45 | 0.50 |
|---|---|---|---|---|---|
| 0.31 | | | | | |
| 0.37 | 1.000 | | | | |
| 0.41 | 1.000 | 1.000 | | | |
| 0.45 | 0.603 | 1.000 | 1.000 | | |
| 0.50 | 0.003 | 0.018 | 0.015 | 1.000 | |

B

| EL | 0.29 | 0.34 | 0.40 | 0.44 | 0.48 |
|---|---|---|---|---|---|
| 0.29 | | | | | |
| 0.34 | 1.000 | | | | |
| 0.40 | 1.000 | 1.000 | | | |
| 0.44 | 1.000 | 1.000 | 1.000 | | |
| 0.48 | 0.071 | 0.015 | 0.004 | 0.861 | |

C

| EL | 0.33 | 0.37 | 0.42 | 0.46 | 0.50 |
|---|---|---|---|---|---|
| 0.33 | | | | | |
| 0.37 | 1.000 | | | | |
| 0.42 | 1.000 | 1.000 | | | |
| 0.46 | 0.194 | 0.256 | 1.000 | | |
| 0.50 | 0.062 | 0.084 | 0.847 | 1.000 | |

**Fig. S8.** The adjusted *P*-values of the *post-hoc* pairwise comparisons Dunn's test for $w^{1118}$ embryos in measurement of *hb* end-to-end distance. (A, B, C) The pairwise comparisons for (A) Em1, (B) Em2 and (C) Em3 respectively. The *P*-values are corrected by Bonferroni correction. The data points with the *P*-values less than 0.05 are marked as red. The AP axis is normalized to embryo length (EL) with 0 and 1 representing the anterior and posterior pole, respectively.

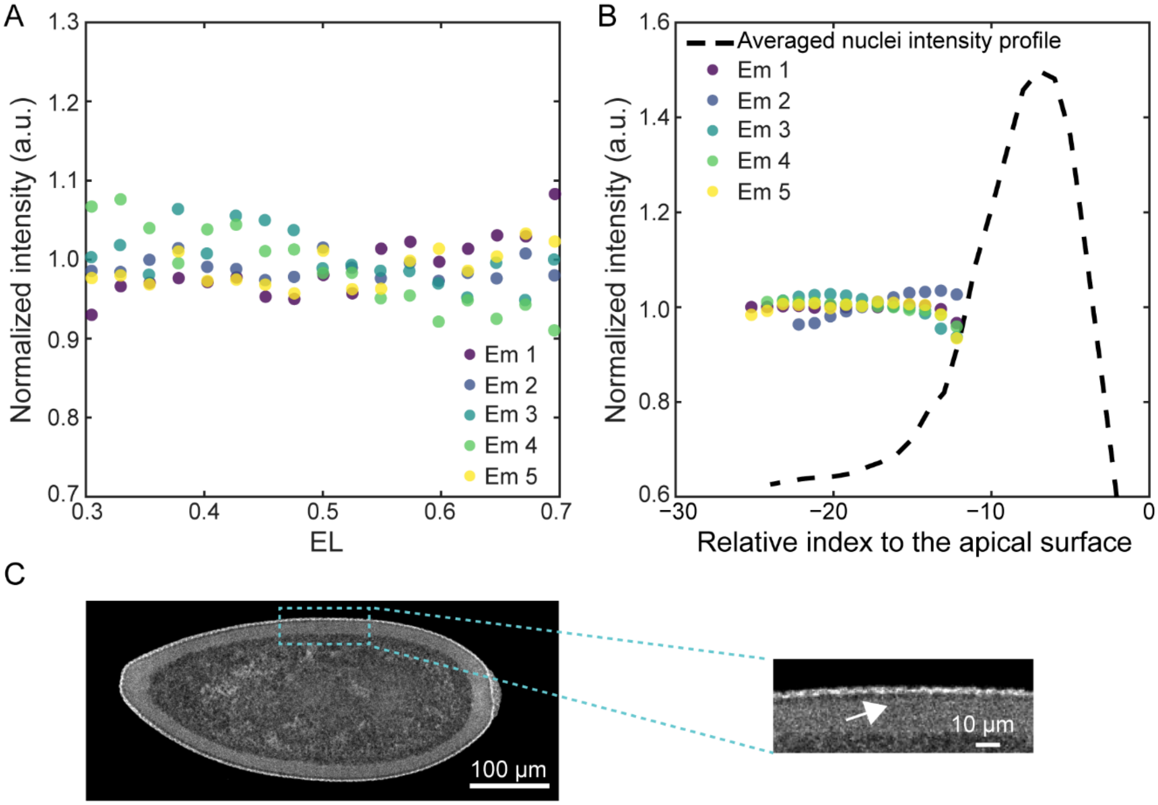


**Fig. S9.** Ribosomal protein RpL10A tagging with GFP is evenly distribution along AP axis and in basal region of *Drosophila* embryo during early nc 14. (A)The normalized fluorescence intensity of RpL10A-GFP along AP axis. The AP axis is normalized to embryo length (EL) with 0 and 1 representing the anterior and posterior pole, respectively. The fluorescence intensity of RpL10A-GFP is first normalized to the co-stained DAPI signals at each AP axis position, then divided by the mean value of the normalized RpL10A-GFP fluorescence intensity in 0.2–0.8 EL for illustration. (B) The normalized fluorescence intensity in the basal region. The fluorescence intensity of RpL10A-GFP is divided by the median value of all intensity values. Nuclei intensity from 5 replicates are averaged and normalized by the normalized fluorescence intensity of RpL10A-GFP for illustration. (C) The representative images of RpL10A-GFP in Drosophila embryo during early nc 14 (Fixed embryo, imaged with GFP fluorescence). Arrow annotates the basal region of cortex. The virgin females of the *$w^{1118}$; P{w[+mC]=GAL4::VP16-nanos.UTR}CG6325[MVD1]* (BDSC4937) were crossed with *w[*]; P{w[+mC]=UAS-GFP-RpL10Ab}BF2* (BDSC42681) from Bloomington *Drosophila* Stock Center for the imaging of the ribosomal protein distribution in the embryos.

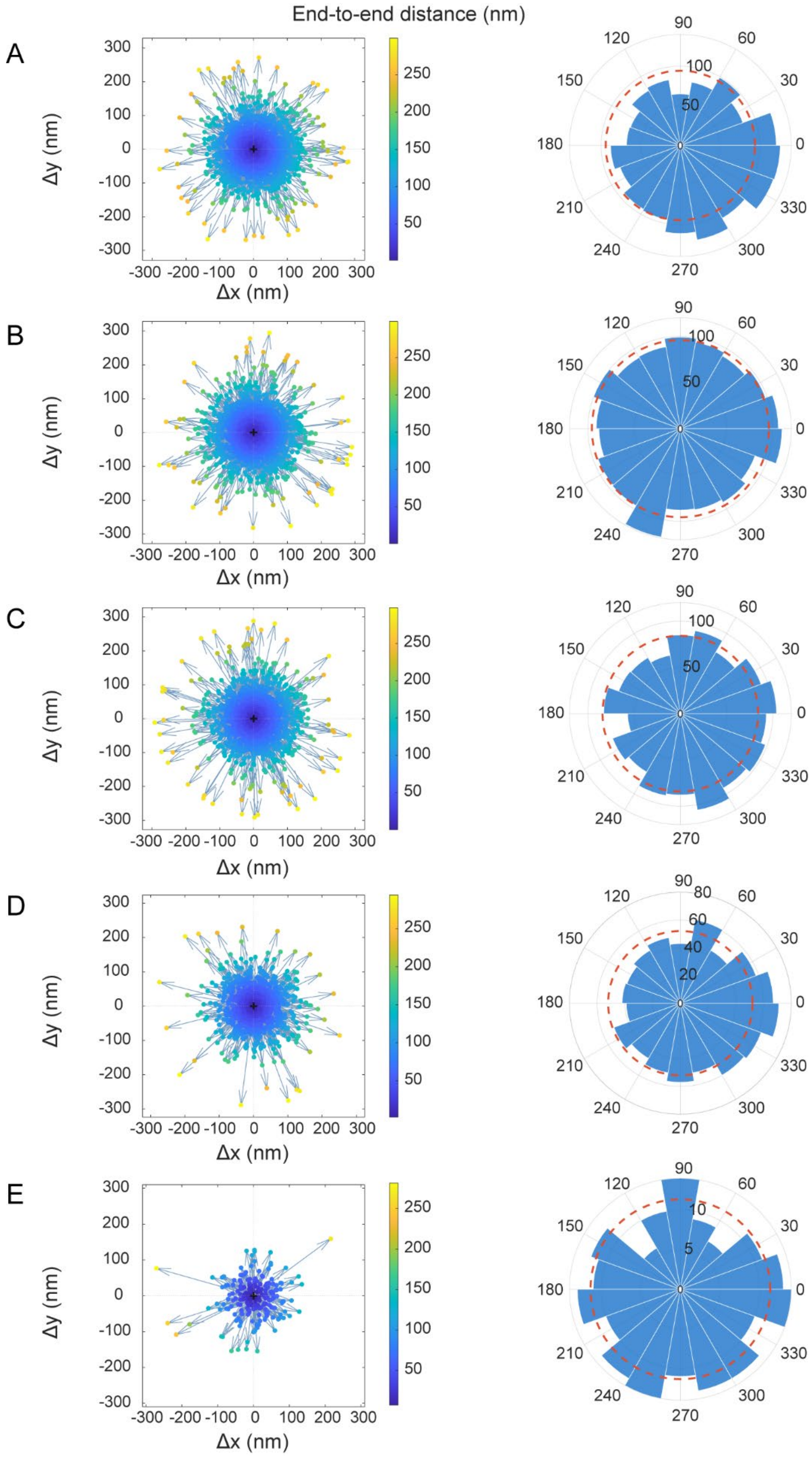
End-to-end distance (nm)
A
B
C
D
E
Δx (nm)
Δy (nm)
90
60
30
0
330
300
270
240
210
180
150
120

**Fig. S10.** Directions and the corresponding count distribution in directions of the AF594-ATTO647N pairs along AP axis. (A–E) The directions (left panel) and count distributions (right panel) from anterior region to boundary region of hb mRNA profile of a representative embryo.

**Table S1.** The sample sizes for permutation-based *KW* test and post-hoc Dunn’s test in *hb-SunTag* replicates

| AP axis positions (EL) | Em1 | Em2 | Em3 |
|---|---|---|---|
| 0.31 | 36 | 30 | 32 |
| 0.33 | 43 | 49 | 41 |
| 0.35 | 56 | 54 | 49 |
| 0.37 | 56 | 56 | 58 |
| 0.39 | 65 | 58 | 65 |
| 0.41 | 72 | 54 | 62 |
| 0.43 | 70 | 56 | 68 |
| 0.45 | 71 | 53 | 70 |
| 0.47 | 64 | 57 | 66 |
| 0.49 | 72 | 57 | 68 |
| 0.51 | 60 | 51 | 64 |
| 0.53 | 51 | 52 | 61 |
| 0.55 | 35 | 55 | 47 |

**Table S2.** The parameter values from model fitting of the mRNA concentration–dependent model to *N*-*TR* data along the AP axis and mRNA count–*TR* data in the basal regions.

| Parameters | Values based on ***N*-*TR*** data | Values based on mRNA count–***TR*** data |
|---|---|---|
| $TR_{\text{offset}}$ | 0.33 ± 0.01 | 0.43 ± 0.05 |
| $TR_{\text{b}}$ | 0.61 ± 0.33 | 0.34 ± 0.03 |
| $C_{\text{T}}$ | 14.10 ± 7.42 | 1214.92 ± 500.13 |

**Table S3.** The parameter values from model fittings of the mRNA concentration–dependent model and constant model to the published *hb* mRNA and Hb protein data in nc 14[1]

| $\boldsymbol{\lambda}$ (min$^{-1}$) | $\boldsymbol{k}_{\mathbf{offset}}$ (min$^{-1}$) | $\boldsymbol{k}_{\mathbf{b}}$ (min$^{-1}$) | $\boldsymbol{C}_{\mathbf{T}}$ | $\boldsymbol{k}$ (min$^{-1}$) |
|---|---|---|---|---|
| 0.005 | 0.046 ± 0.004 | 0.223 ± 0.046 | 35.990 ± 7.892 | 0.038 ± 0.004 |
| 0.010 | 0.053 ± 0.005 | 0.205 ± 0.046 | 37.835 ± 9.515 | 0.045 ± 0.004 |
| 0.015 | 0.060 ± 0.006 | 0.189 ± 0.047 | 40.084 ± 11.738 | 0.051 ± 0.004 |
| 0.020 | 0.067 ± 0.007 | 0.173 ± 0.047 | 42.870 ± 14.848 | 0.057 ± 0.004 |
| 0.025 | 0.074 ± 0.009 | 0.157 ± 0.046 | 46.395 ± 19.339 | 0.063 ± 0.003 |

**Table S4.** Probe sequences for *SunTag* smFISH

| Index | Sequences |
|---|---|
| 24× SunTag_1 | ccacttcgttctcaagatga |
| 24× SunTag_2 | gccagaacctttcttaagac |
| 24× SunTag_3 | ttgaaagcagttcttctcca |
| 24× SunTag_4 | ctcattttccaggtggtaat |
| 24× SunTag_5 | aatttttgctcagcaactcc |
| 24× SunTag_6 | ttctttagtcgtgctacttc |
| 24× SunTag_7 | tttcgagagtaactcctcac |
| 24× SunTag_8 | ccacttcgttttcgagatga |
| 24× SunTag_9 | gataatagctcttctccaga |
| 24× SunTag_10 | tcattttcgaggtggtagtt |
| 24× SunTag_11 | acttcccttttttaagcgtg |
| 24× SunTag_12 | tcttggatagtagctcttca |
| 24× SunTag_13 | ctacctcgttctcaagatga |
| 24× SunTag_14 | tagttcttcgagagcagttc |
| 24× SunTag_15 | gatcccttttttaatcgagc |
| 24× SunTag_16 | tgaaagtagttcctcaccac |
| 24× SunTag_17 | cttcgttttcgaggtggtaa |
| 24× SunTag_18 | ccctgaacctttctttaatc |
| 24× SunTag_19 | tactcagtaattcttcaccc |
| 24× SunTag_20 | ctacctcattttccagatga |
| 24× SunTag_21 | tttcgatagcaactcttcgc |
| 24× SunTag_22 | tttttgagcctagcaacttc |
| 24× SunTag_23 | agtggtagtttttcgagagc |
| 24× SunTag_24 | tttgctcaataactcctcgc |
| 24× SunTag_25 | cgcgacttcgttctctaaat |
| 24× SunTag_26 | ttcgataagagttcttcgcc |
| 24× SunTag_27 | Ctcattttcgaggtggtagt |
| 24× SunTag_28 | agtggtagttcttgctcaag |
| 24× SunTag_29 | attcttgctgagcaattcct |
| 24× SunTag_30 | cgacttcgttctccaaatga |
| 24× SunTag_31 | tttactcaacaattcctccc |
| 24× SunTag_32 | cgacttcattttccaagtgg |
| 24× SunTag_33 | ttgctcaataactcttcgcc |
| 24× SunTag_34 | ttcgttctccaagtggtaat |

| | |
|---|---|
| 24× SunTag_35 | agttcttcgataagagctcc |
| 24× SunTag_36 | gcgacttcattctctaagtg |
| 24× SunTag_37 | agtggtagttcttgctcaag |
| 24× SunTag_38 | ttagatagtaactcttcccc |
| 24× SunTag_39 | cctcgttctcgagatgataa |
| 24× SunTag_40 | gatagttcttcgacaggagt |
| 24× SunTag_41 | cctttttaagtcttgcaacc |
| 24× SunTag_42 | ttcttactgagtagttcctc |
| 24× SunTag_43 | tcctgatcctttcttcaaac |
| 24× SunTag_44 | cttttgagagcagttcttcg |
| 24× SunTag_45 | gcaacctcattttccaaatg |
| 24× SunTag_46 | tgccacttcccttttttaaa |
| 24× SunTag_47 | tttcgacagaagttcctcac |
| 24× SunTag_48 | taagtcgggctacttcattc |

**Table S5.** Probe sequences for *hb* smFISH (*hb* mRNA and Hb protein co-staining)

| Index | Sequences |
|---|---|
| *hb*_1 | atcttggcggctctagacgg |
| *hb*_2 | ggctgtcgtctcccagttct |
| *hb*_3 | tgtgctgctcgtagttggtc |
| *hb*_4 | cgcgaacatgctgttgtacc |
| *hb*_5 | gacctggctcctgtttgata |
| *hb*_6 | acgctattcccgtcgagatg |
| *hb*_7 | atgggcgattggcgcggact |
| *hb*_8 | ttccaggtgattggtcgagg |
| *hb*_9 | atgggttgctgctgaagctg |
| *hb*_10 | ggtcatggcgcacagggtat |
| *hb*_11 | gatcgttttggctgggtgat |
| *hb*_12 | catcgtaatgctgcaggctg |
| *hb*_13 | cccatcagatggtgatggtg |
| *hb*_14 | tggcgtcagcggattgaatc |
| *hb*_15 | gcatgggattgggcagacca |
| *hb*_16 | gcagattgccgccatagaag |
| *hb*_17 | atgtgggcgtgggctgcgga |
| *hb*_18 | actgcaacgggcgcaattgt |
| *hb*_19 | acttctcgctgctgccagtg |
| *hb*_20 | atgggtggtgttagtgcctg |
| *hb*_21 | cgacttaggcggtgtgacat |
| *hb*_22 | tgctccttctccggctcaat |
| *hb*_23 | cggccatgtacttcatgtcc |
| *hb*_24 | ttggtatcatcgtcctcgga |
| *hb*_25 | attgtagatgggcatccgga |
| *hb*_26 | agttcttcatcttgccgtgc |
| *hb*_27 | aagtccaccttggtgatggc |
| *hb*_28 | tcggtgacgaacgggcactt |
| *hb*_29 | ggtactccaagtggtgcttg |
| *hb*_30 | tggttcttgtgcttccggat |
| *hb*_31 | tgttgacacacgtgtagctg |
| *hb*_32 | cggtgcgagtttagcatgga |
| *hb*_33 | gatacacagaactgtgcgac |
| *hb*_34 | tcacaatccgcacaacggta |
| *hb*_35 | gtggcaatacttggtggcgt |

| | |
|---|---|
| *hb*_36 | tgcgcagatgcagcttgaag |
| *hb*_37 | atgccgggcttgtgaccata |
| *hb*_38 | ggtgccatcctcgtccaaaa |
| *hb*_39 | cgatgaccaacgagggattc |
| *hb*_40 | accacgacgcgtgccgtaaa |
| *hb*_41 | cggtccaccattcttgctct |
| *hb*_42 | tgccacttcctccactggca |
| *hb*_43 | acatttgacttccggctgcc |
| *hb*_44 | ttgctgcggagcgacagctg |
| *hb*_45 | gctgagctggctgagattgc |
| *hb*_46 | actcagctgagatgtggcga |

**Table S6.** smiFISH probes for targeting 5′ end of *hb* mRNA

| Index | Sequences |
|---|---|
| *hb*_5_1_i | CCTCCTAAGTTTCGAGCTGGACTCAGTGcgcagtggtaacaaactcgt |
| *hb*_5_2_i | CCTCCTAAGTTTCGAGCTGGACTCAGTGtattggtgctgcttctgttg |
| *hb*_5_3_i | CCTCCTAAGTTTCGAGCTGGACTCAGTGttcgtaaggatttgcaagta |
| *hb*_5_4_i | CCTCCTAAGTTTCGAGCTGGACTCAGTGatattccaaatttgtcggga |
| *hb*_5_5_i | CCTCCTAAGTTTCGAGCTGGACTCAGTGcagtgcgtatgattgcgatt |
| *hb*_5_6_i | CCTCCTAAGTTTCGAGCTGGACTCAGTGcgaacatgctgttgtaccag |
| *hb*_5_7_i | CCTCCTAAGTTTCGAGCTGGACTCAGTGtggctcctgtttgatatttg |
| *hb*_5_8_i | CCTCCTAAGTTTCGAGCTGGACTCAGTGtattcccgtcgagatgatga |
| *hb*_5_9_i | CCTCCTAAGTTTCGAGCTGGACTCAGTGaggtgattggtcgagggaat |
| *hb*_5_10_i | CCTCCTAAGTTTCGAGCTGGACTCAGTGtgttttgatcgttttggctg |
| *hb*_5_11_i | CCTCCTAAGTTTCGAGCTGGACTCAGTGttagcatcgtaatgctgcag |
| *hb*_5_12_i | CCTCCTAAGTTTCGAGCTGGACTCAGTGcgtcagcggattgaatccac |
| *hb*_5_13_i | CCTCCTAAGTTTCGAGCTGGACTCAGTGatagaagtgctgcatgggat |
| *hb*_5_14_i | CCTCCTAAGTTTCGAGCTGGACTCAGTGaattgtggaggcagatgtgg |
| *hb*_5_15_i | CCTCCTAAGTTTCGAGCTGGACTCAGTGtgttagtgcctgcaacttct |
| *hb*_5_16_i | CCTCCTAAGTTTCGAGCTGGACTCAGTGgacttaggcggtgtgacatc |
| *hb*_5_17_i | CCTCCTAAGTTTCGAGCTGGACTCAGTGattcgactgactcgacttgg |
| *hb*_5_18_i | CCTCCTAAGTTTCGAGCTGGACTCAGTGgagttcgacatctgatcgtg |
| *hb*_5_19_i | CCTCCTAAGTTTCGAGCTGGACTCAGTGcatgtacttcatgtcctcgc |
| *hb*_5_20_i | CCTCCTAAGTTTCGAGCTGGACTCAGTGtatcatcgtcctcggactcg |
| *hb*_5_21_i | CCTCCTAAGTTTCGAGCTGGACTCAGTGtagatgggcatccggatgtt |
| *hb*_5_22_i | CCTCCTAAGTTTCGAGCTGGACTCAGTGcttcatcttgccgtgcgaat |
| *hb*_5_23_i | CCTCCTAAGTTTCGAGCTGGACTCAGTGcgcaggtcttgcacttgtag |
| *hb*_5_24_i | CCTCCTAAGTTTCGAGCTGGACTCAGTGagaagtccaccttggtgatg |
| *hb*_5_25_i | CCTCCTAAGTTTCGAGCTGGACTCAGTGatcttgtctggtttcatgtg |
| AF594 FLAPX | CACTGAGTCCAGCTCGAAACTTAGGAGG |

**Table S7.** smiFISH probes for targeting 3′ end of *hb* mRNA

| Index | Sequences |
|---|---|
| *hb*_3_1_i | TTACACTCGGACCTCGTCGACATGCATTtcgaactggcactggtattg |
| *hb*_3_2_i | TTACACTCGGACCTCGTCGACATGCATTcagtacttgcactcgtagat |
| *hb*_3_3_i | TTACACTCGGACCTCGTCGACATGCATTggcgtccttgaagaagatat |
| *hb*_3_4_i | TTACACTCGGACCTCGTCGACATGCATTccatgtgaatggtgtagagc |
| *hb*_3_5_i | TTACACTCGGACCTCGTCGACATGCATTtgcacttgaacacatcgtcg |
| *hb*_3_6_i | TTACACTCGGACCTCGTCGACATGCATTatgtgaacgaagaggccgac |
| *hb*_3_7_i | TTACACTCGGACCTCGTCGACATGCATTcttaggagtgagcattcctg |
| *hb*_3_8_i | TTACACTCGGACCTCGTCGACATGCATTacaaggtgatggtgatgggg |
| *hb*_3_9_i | TTACACTCGGACCTCGTCGACATGCATTggcttatgtacaattttcga |
| *hb*_3_10_i | TTACACTCGGACCTCGTCGACATGCATTtaggtctagaattagcggct |
| *hb*_3_11_i | TTACACTCGGACCTCGTCGACATGCATTggtggctacagttcaataca |
| *hb*_3_12_i | TTACACTCGGACCTCGTCGACATGCATTgtgtataggagacagattga |
| *hb*_3_13_i | TTACACTCGGACCTCGTCGACATGCATTgttttcagggtttttagtcg |
| *hb*_3_14_i | TTACACTCGGACCTCGTCGACATGCATTcttatgtttctccatgcatt |
| *hb*_3_15_i | TTACACTCGGACCTCGTCGACATGCATTgttgtttcgcttgactcaac |
| *hb*_3_16_i | TTACACTCGGACCTCGTCGACATGCATTtttgactttggactgttggt |
| *hb*_3_17_i | TTACACTCGGACCTCGTCGACATGCATTagaactgagtgttatgcgca |
| *hb*_3_18_i | TTACACTCGGACCTCGTCGACATGCATTgtcattattgccaaaaggtt |
| *hb*_3_19_i | TTACACTCGGACCTCGTCGACATGCATTtcttctttcgtcagtttcag |
| *hb*_3_20_i | TTACACTCGGACCTCGTCGACATGCATTcgcatcttagctactcttta |
| *hb*_3_21_i | TTACACTCGGACCTCGTCGACATGCATTaattttgatccgttgctcag |
| *hb*_3_22_i | TTACACTCGGACCTCGTCGACATGCATTcgacttagattttatggggt |
| *hb*_3_23_i | TTACACTCGGACCTCGTCGACATGCATTgtctcgaaattcgtttcatg |
| *hb*_3_24_i | TTACACTCGGACCTCGTCGACATGCATTatcaaggattacactgggct |
| *hb*_3_25_i | TTACACTCGGACCTCGTCGACATGCATTcatttcgtgggcaaatatct |
| ATTO 647N FLAPY | AATGCATGTCGACGAGGTCCGAGTGTAA |

**Table S8.** smiFISH probes targeting *hb* CDS for *hb-def* embryo selection

| Index | Sequences |
|---|---|
| *hb*_CDSi_1 | TTACACTCGGACCTCGTCGACATGCATTtgtgctgctcgtagttggtc |
| *hb*_CDSi_2 | TTACACTCGGACCTCGTCGACATGCATTcgcgaacatgctgttgtacc |
| *hb*_CDSi_3 | TTACACTCGGACCTCGTCGACATGCATTgacctggctcctgtttgata |
| *hb*_CDSi_4 | TTACACTCGGACCTCGTCGACATGCATTacgctattcccgtcgagatg |
| *hb*_CDSi_5 | TTACACTCGGACCTCGTCGACATGCATTatgggcgattggcgcggact |
| *hb*_CDSi_6 | TTACACTCGGACCTCGTCGACATGCATTttccaggtgattggtcgagg |
| *hb*_CDSi_7 | TTACACTCGGACCTCGTCGACATGCATTatgggttgctgctgaagctg |
| *hb*_CDSi_8 | TTACACTCGGACCTCGTCGACATGCATTggtcatggcgcacagggtat |
| *hb*_CDSi_9 | TTACACTCGGACCTCGTCGACATGCATTgatcgttttggctgggtgat |
| *hb*_CDSi_10 | TTACACTCGGACCTCGTCGACATGCATTcatcgtaatgctgcaggctg |
| *hb*_CDSi_11 | TTACACTCGGACCTCGTCGACATGCATTcccatcagatggtgatggtg |
| *hb*_CDSi_12 | TTACACTCGGACCTCGTCGACATGCATTtggcgtcagcggattgaatc |
| *hb*_CDSi_13 | TTACACTCGGACCTCGTCGACATGCATTgcatgggattgggcagacca |
| *hb*_CDSi_14 | TTACACTCGGACCTCGTCGACATGCATTgcagattgccgccatagaag |
| *hb*_CDSi_15 | TTACACTCGGACCTCGTCGACATGCATTatgtgggcgtgggctgcgga |
| *hb*_CDSi_16 | TTACACTCGGACCTCGTCGACATGCATTactgcaacgggcgcaattgt |
| *hb*_CDSi_17 | TTACACTCGGACCTCGTCGACATGCATTacttctcgctgctgccagtg |
| *hb*_CDSi_18 | TTACACTCGGACCTCGTCGACATGCATTatgggtggtgttagtgcctg |
| *hb*_CDSi_19 | TTACACTCGGACCTCGTCGACATGCATTcgacttaggcggtgtgacat |
| *hb*_CDSi_20 | TTACACTCGGACCTCGTCGACATGCATTtgctccttctccggctcaat |
| *hb*_CDSi_21 | TTACACTCGGACCTCGTCGACATGCATTcggccatgtacttcatgtcc |
| *hb*_CDSi_22 | TTACACTCGGACCTCGTCGACATGCATTttggtatcatcgtcctcgga |
| *hb*_CDSi_23 | TTACACTCGGACCTCGTCGACATGCATTattgtagatgggcatccgga |
| *hb*_CDSi_24 | TTACACTCGGACCTCGTCGACATGCATTagttcttcatcttgccgtgc |
| *hb*_CDSi_25 | TTACACTCGGACCTCGTCGACATGCATTaagtccaccttggtgatggc |
| *hb*_CDSi_26 | TTACACTCGGACCTCGTCGACATGCATTtcggtgacgaacgggcactt |
| *hb*_CDSi_27 | TTACACTCGGACCTCGTCGACATGCATTggtactccaagtggtgcttg |
| *hb*_CDSi_28 | TTACACTCGGACCTCGTCGACATGCATTtggttcttgtgcttccggat |
| *hb*_CDSi_29 | TTACACTCGGACCTCGTCGACATGCATTtgttgacacacgtgtagctg |
| *hb*_CDSi_30 | TTACACTCGGACCTCGTCGACATGCATTcggtgcgagtttagcatgga |
| *hb*_CDSi_31 | TTACACTCGGACCTCGTCGACATGCATTgatacacagaactgtgcgac |
| *hb*_CDSi_32 | TTACACTCGGACCTCGTCGACATGCATTtcacaatccgcacaacggta |
| *hb*_CDSi_33 | TTACACTCGGACCTCGTCGACATGCATTgtggcaatacttggtggcgt |
| *hb*_CDSi_34 | TTACACTCGGACCTCGTCGACATGCATTtgcgcagatgcagcttgaag |
| *hb*_CDSi_35 | TTACACTCGGACCTCGTCGACATGCATTatgccgggcttgtgaccata |

| | |
|---|---|
| *hb*_CDSi_36 | TTACACTCGGACCTCGTCGACATGCATTggtgccatcctcgtccaaaa |
| *hb*_CDSi_37 | TTACACTCGGACCTCGTCGACATGCATTcgatgaccaacgagggattc |
| *hb*_CDSi_38 | TTACACTCGGACCTCGTCGACATGCATTaccacgacgcgtgccgtaaa |
| *hb*_CDSi_39 | TTACACTCGGACCTCGTCGACATGCATTcggtccaccattcttgctct |
| *hb*_CDSi_40 | TTACACTCGGACCTCGTCGACATGCATTtgccacttcctccactggca |
| *hb*_CDSi_41 | TTACACTCGGACCTCGTCGACATGCATTacatttgacttccggctgcc |
| *hb*_CDSi_42 | TTACACTCGGACCTCGTCGACATGCATTttgctgcggagcgacagctg |
| *hb*_CDSi_43 | TTACACTCGGACCTCGTCGACATGCATTgctgagctggctgagattgc |
| *hb*_CDSi_44 | TTACACTCGGACCTCGTCGACATGCATTactcagctgagatgtggcga |
| AF488_FALPY | AATGCATGTCGACGAGGTCCGAGTGTAA |

**Table S9.** smiFISH probes for dual-label co-localization control

| Index | Sequences |
|---|---|
| hb_loc_even1 | CCTCCTAAGTTTCGAGCTGGACTCAGTGtattggtgctgcttctgttg |
| hb_loc_even2 | CCTCCTAAGTTTCGAGCTGGACTCAGTGatattccaaatttgtcggga |
| hb_loc_even3 | CCTCCTAAGTTTCGAGCTGGACTCAGTGcgaacatgctgttgtaccag |
| hb_loc_even4 | CCTCCTAAGTTTCGAGCTGGACTCAGTGtattcccgtcgagatgatga |
| hb_loc_even5 | CCTCCTAAGTTTCGAGCTGGACTCAGTGtgttttgatcgttttggctg |
| hb_loc_even6 | CCTCCTAAGTTTCGAGCTGGACTCAGTGcgtcagcggattgaatccac |
| hb_loc_even7 | CCTCCTAAGTTTCGAGCTGGACTCAGTGtgttagtgcctgcaacttct |
| hb_loc_even8 | CCTCCTAAGTTTCGAGCTGGACTCAGTGattcgactgactcgacttgg |
| hb_loc_even9 | CCTCCTAAGTTTCGAGCTGGACTCAGTGactcggccatgtacttcatg |
| hb_loc_even10 | CCTCCTAAGTTTCGAGCTGGACTCAGTGtgccgtgcgaattgtagatg |
| hb_loc_even11 | CCTCCTAAGTTTCGAGCTGGACTCAGTGagaagtccaccttggtgatg |
| hb_loc_even12 | CCTCCTAAGTTTCGAGCTGGACTCAGTGtgaactcggtgacgaacggg |
| hb_loc_even13 | CCTCCTAAGTTTCGAGCTGGACTCAGTGcttttggttcttgtgcttcc |
| hb_loc_even14 | CCTCCTAAGTTTCGAGCTGGACTCAGTGggatttgttgacacacgtgt |
| hb_loc_even15 | CCTCCTAAGTTTCGAGCTGGACTCAGTGcggtactgatacacagaact |
| hb_loc_even16 | CCTCCTAAGTTTCGAGCTGGACTCAGTGagcttgaagctgtggcaata |
| hb_loc_even17 | CCTCCTAAGTTTCGAGCTGGACTCAGTGcatcctcgtccaaaaccatg |
| hb_loc_even18 | CCTCCTAAGTTTCGAGCTGGACTCAGTGacgacgcgtgccgtaaacat |
| hb_loc_even19 | CCTCCTAAGTTTCGAGCTGGACTCAGTGgacagctgcaacatttgact |
| hb_loc_even20 | CCTCCTAAGTTTCGAGCTGGACTCAGTGcagctgagatgtggcgactg |
| hb_loc_even21 | CCTCCTAAGTTTCGAGCTGGACTCAGTGagcatctggagattgaggtt |
| hb_loc_even22 | CCTCCTAAGTTTCGAGCTGGACTCAGTGtggttctgttgctgcagttg |
| hb_loc_even23 | CCTCCTAAGTTTCGAGCTGGACTCAGTGattcgagctgctcatcagag |
| hb_loc_even24 | CCTCCTAAGTTTCGAGCTGGACTCAGTGtttcagttgctcgggagatg |
| hb_loc_odd1 | TTACACTCGGACCTCGTCGACATGCATTcgcagtggtaacaaactcgt |
| hb_loc_odd2 | TTACACTCGGACCTCGTCGACATGCATTttcgtaaggatttgcaagta |
| hb_loc_odd3 | TTACACTCGGACCTCGTCGACATGCATTcagtgcgtatgattgcgatt |
| hb_loc_odd4 | TTACACTCGGACCTCGTCGACATGCATTtggctcctgtttgatatttg |
| hb_loc_odd5 | TTACACTCGGACCTCGTCGACATGCATTgtgattggtcgagggaatgg |
| hb_loc_odd6 | TTACACTCGGACCTCGTCGACATGCATTttagcatcgtaatgctgcag |
| hb_loc_odd7 | TTACACTCGGACCTCGTCGACATGCATTatagaagtgctgcatgggat |
| hb_loc_odd8 | TTACACTCGGACCTCGTCGACATGCATTgacttaggcggtgtgacatc |
| hb_loc_odd9 | TTACACTCGGACCTCGTCGACATGCATTctggagttcgacatctgatc |
| hb_loc_odd10 | TTACACTCGGACCTCGTCGACATGCATTcggatgttggtatcatcgtc |
| hb_loc_odd11 | TTACACTCGGACCTCGTCGACATGCATTtcttgcacttgtagttcttc |

| | |
|---|---|
| hb_loc_odd12 | TTACACTCGGACCTCGTCGACATGCATTatcttgtctggtttcatgtg |
| hb_loc_odd13 | TTACACTCGGACCTCGTCGACATGCATTatatggtactccaagtggtg |
| hb_loc_odd14 | TTACACTCGGACCTCGTCGACATGCATTtgcatttgtcgcactggaag |
| hb_loc_odd15 | TTACACTCGGACCTCGTCGACATGCATTgacttgcggtgcgagtttag |
| hb_loc_odd16 | TTACACTCGGACCTCGTCGACATGCATTtggtggcgtaatcacaatcc |
| hb_loc_odd17 | TTACACTCGGACCTCGTCGACATGCATTttgtgaccatacttgcgcag |
| hb_loc_odd18 | TTACACTCGGACCTCGTCGACATGCATTatgaccaacgagggattcgg |
| hb_loc_odd19 | TTACACTCGGACCTCGTCGACATGCATTcaatcggtccaccattcttg |
| hb_loc_odd20 | TTACACTCGGACCTCGTCGACATGCATTtgagctggctgagattgctg |
| hb_loc_odd21 | TTACACTCGGACCTCGTCGACATGCATTcacactcttggcaggagagg |
| hb_loc_odd22 | TTACACTCGGACCTCGTCGACATGCATTcgcattcttggcgacaattg |
| hb_loc_odd23 | TTACACTCGGACCTCGTCGACATGCATTgcagagtccactgacttacg |
| hb_loc_odd24 | TTACACTCGGACCTCGTCGACATGCATTatcgcagttgtaacagggtg |
